# Salinity adaptation in a copepod reveals contrasting genomic architectures and the limits of genomic offsets

**DOI:** 10.64898/2026.11.13.766083

**Authors:** Alexandra Hahn, Jennifer C. Nascimento-Schulze, Georgia Avgerinou, Lara Schmittmann, Julian Rethemeier, Willi Rath, Reid S. Brennan

**Affiliations:** Research Division Marine Ecology – Evolutionary Ecology, GEOMAR Helmholtz Centre for Ocean Research Kiel, Kiel, Germany; Thünen-Institute of Fisheries Ecology, Bremerhaven, Germany; Research Division Ocean Circulation and Climate Dynamics – Ocean Dynamics, GEOMAR Helmholtz Centre for Ocean Research Kiel, Kiel, Germany; Kiel University (CAU), Kiel, Germany; Marine Mammal and Turtle Division, Southeast Fisheries Science Center, National Marine Fisheries Service, National Oceanic and Atmospheric Administration, Miami FL, USA

**Keywords:** local adaptation, *Acartia tonsa*, Baltic Sea, osmoregulation, polygenic adaptation, genomic architecture, genomic offset, salinity tolerance, resilience, common-garden experiments, population genomics

## Abstract

Gene flow is expected to homogenize populations, yet adaptive divergence can persist across environmental gradients. We investigated how genomic architectures and physiological tolerance shape salinity adaptation and resilience by combining whole-genome pool-sequencing of 11 copepod populations (*Acartia tonsa)* with common-garden physiology and dispersal modelling. Divergence between the high-salinity North Sea and low-salinity Baltic Sea was driven by few strongly differentiated genomic regions, while adaptation along the Baltic salinity gradient persisted despite gene flow and was highly polygenic, with candidate loci enriched for osmoregulatory processes. In common gardens, low-salinity populations survived reduced salinity better, corroborating genomic signals. Genomic offsets failed to capture range restrictions predicted by physiology-informed distribution models, likely because salinity tolerance behaves as a threshold trait, most impactful at range margins. Accounting for local adaptation reduced predicted habitat loss by 79%. Our results demonstrate that integrating physiological thresholds with genomics can improve forecasts of species’ responses to environmental change.

## 1 INTRODUCTION

Environmental gradients are powerful drivers of natural selection, shaping patterns of adaptation and divergence across space (Sanford & Kelly, 2011). Yet the extent to which populations can adapt to local environments is strongly variable and dependent on a complex interaction of factors including trait architecture, strength of selection, gene flow, and genetic drift. In systems where gene flow is high, strong selection may favor few and tightly linked loci, as clustered architectures are more resistant to disruptions by recombination under gene flow (Yeaman & Whitlock, 2011). However, this expectation rests on the assumption that adaptation is driven by few large-effect loci, each individually targeted by selection. In cases where adaptation is polygenic and many genes underlie a trait, no single locus is the main target of selection and adaptation might persist despite gene flow (Yeaman, 2015), especially if genetic redundancy is high (Barghi et al., 2020). This distinction between genomic architectures has important implications for which genomic signatures we expect to observe, and for our understanding how local adaptation can be maintained even in connected populations.

Local adaptation arises when populations evolve heritable fitness advantages in their local environment relative to individuals from other populations (Kawecki & Ebert, 2004) and therefore is one of the main mechanisms that allows species persistence across heterogeneous environments. Studying local adaptation is of particular interest in the context of global change, as populations currently adapted to different environments can provide a window into evolutionary responses to shifting conditions over time, i.e., space-for-time substitutions (Lovell et al., 2023). The underlying assumption is that the standing genetic variation driving local adaptation along contemporary gradients can also facilitate temporal adaptation as those gradients shift (Walters & Berger, 2019). In this context, genomic offsets have emerged as a tool for forecasting population vulnerability to future conditions by quantifying the mismatch between current and predicted future allele frequencies (Fitzpatrick & Keller, 2015). However, offset predictions rely on key assumptions that require careful validation, including that genotype-environment associations reflect current adaptive optima and that larger offsets correlate with larger fitness declines (Rellstab et al., 2021; Ahrens et al., 2025). Linking genomic signals to phenotypic fitness data from common-garden experiments can be used for validation and to obtain a more holistic understanding of the mechanisms and consequences of adaptation.

Planktonic organisms present a particularly compelling opportunity to understand adaptation with gene flow in marine systems. Many planktonic species are characterized by large population sizes and short generation times, traits that could facilitate rapid evolutionary responses. Indeed, planktonic copepods have been shown to rapidly adapt to various conditions over relatively few generations (Posavi et al., 2020; Dam et al., 2021; Brennan et al., 2022; Stern et al., 2022). However, high dispersal of planktonic organisms through ocean currents has traditionally been assumed to limit population differentiation and local adaptation in pelagic systems (Haldane, 1930; Tigano & Friesen, 2016). In recent years, a growing body of evidence suggests that local adaptation in marine systems is more common than previously thought, even in planktonic species (Sanford & Kelly, 2011). For example, in highly connected populations of planktonic diatoms, fine-scale population structure and signatures of selection have been found along an environmental gradient (Pinseel et al., 2025). Yet for most planktonic taxa, we still lack understanding of how adaptive variation is structured across seascapes. In this study, we leverage the planktonic copepod *Acartia tonsa* as a model to understand the role of genomic architecture in maintaining local adaptation despite gene flow, as well as predicting the species’ resilience to future change. This species is known for its broad salinity tolerance (Cervetto et al., 1999) and as an integral part of zooplankton communities worldwide where it serves an important component in the diet of numerous fish species (Turner, 1984; Mauchline, 1998; Höper et al., 2025). In addition, short generation times on the order of weeks (Leandro et al., 2006), and the possibility of establishing long term experimental cultures and common gardens, makes *A. tonsa* an ideal model organism for studying adaptation and resilience to global change (Dam, 2013).

Salinity is one of the most important abiotic stressors for marine organisms and a main driver of species distribution (Lee et al., 2022). For most marine species, transitioning between salinities is extremely challenging and energetically costly, as osmotic and ionic regulation demands coordinated responses across ion transport, cell volume regulation, and signalling (Mantel & Farmer, 1983). As a result, salinity tolerance is a complex trait and the genomic architecture of salinity adaptation is often polygenic, with adaptive signals distributed across the genome (Brennan et al., 2018; Lee, 2023). Alternatively, co-adapted alleles may be captured and maintained by structural variants such as chromosomal inversions, which suppress recombination and protect locally adapted haplotypes from disruption by gene flow, facilitating adaptation across environmental gradients (Kirkpatrick & Barton, 2006; Wellenreuther et al., 2019). The Baltic Sea is an ideal natural laboratory for studying these dynamics. It spans one of the most extreme salinity gradients worldwide, from fully marine in the North Sea transition area to almost freshwater in the north-eastern basins (Snoeijs-Leijonmalm et al., 2017). Distinct genetic clines along this gradient are well documented, and in several species, including seaweed flies, plaice, and cod, chromosomal inversions have been directly linked to environmental clines from the North Sea to the Baltic and appear to play an important role in adaptation (Day et al., 1983; Johannesson et al., 2020; Le Moan et al., 2021; Matschiner et al., 2022).

Understanding the genomic architecture of salinity adaptation in such systems is relevant to predict biological responses to future change. Salinity regimes are projected to shift substantially under global change (Durack et al., 2012; Du et al., 2019), and the Baltic has already experienced local salinity reductions of up to 1.14 PSU over the past 60 years (Kankaanpää et al., 2023), and models predict ongoing freshening under climate change (Kniebusch et al., 2019). Therefore, the Baltic Sea serves as a model for understanding local adaptation along steep environmental gradients, but is also considered a ‘time machine’ as it already poses some of the evolutionary challenges that species will face as salinity regimes shift globally (Reusch et al., 2018).

The aim of this study was to understand salinity adaptation across different spatial scales and predict population resilience to changes in salinity. We used whole-genome-resequencing (WGR) data generated from 11 pools of *A. tonsa* copepods sampled along the salinity cline throughout North and Baltic Sea to identify genetic regions underlying salinity adaptation and combined this genomic data set with common-garden experiments on 10 lab populations originating from the same sampling locations (Figure 1A and B). We further integrated Lagrangian dispersal simulations to assess how ocean currents may affect populations connectivity and spatial patterns of adaptation. Lastly, we compared predictions of resilience from genomic vulnerability versus a newly derived physiological model that incorporated experimentally derived survival estimates. Genomic predictions of vulnerability performed poorly for this system and instead the physiologically derived model more accurately reflected the biology of this species. Importantly, allowing for adaptive potential in the physiological model significantly reduced the predictions of habitat loss in the Baltic Sea in the future. By integrating seascape genomics, experimental fitness, and dispersal simulations across a strong environmental gradient, our study is an interdisciplinary assessment of local adaptation and resilience in a highly dispersive marine species. Our results illustrate how planktonic species can serve as powerful models for understanding local adaptation and species’ responses to ongoing environmental change.

**Figure 1.**
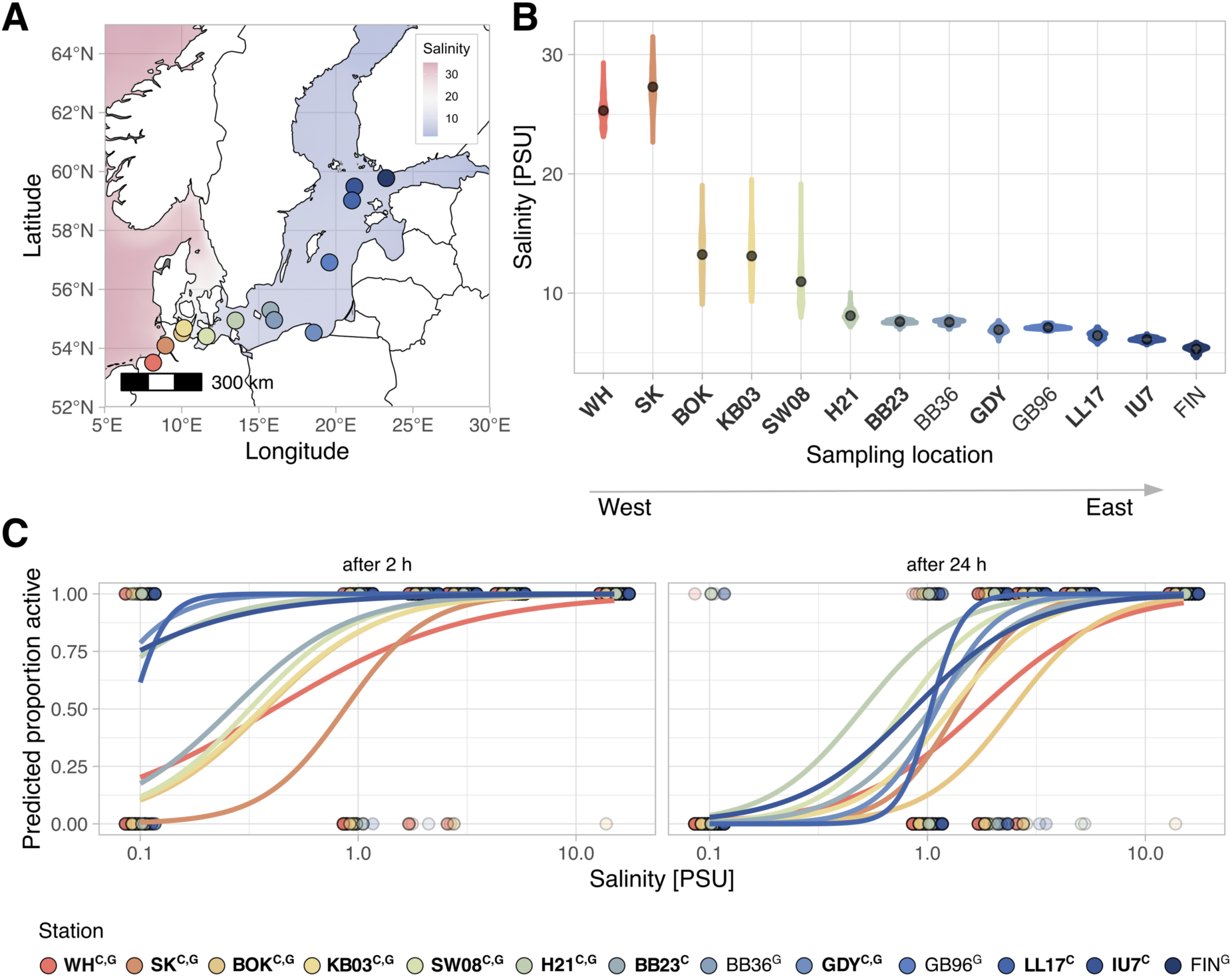
Sampling sites and survival to low salinity, superscript letters in Figure legend indicate data type available for this location (C = culture, G = genomic). A) Map of all sampling locations for this study, colors of sampling locations correspond to Table 1 and are consistent throughout the manuscript. B) Surface salinity per sampling locations ordered from West to East, data from CMEMS for daily means from June-October for sampling year and five years prior, locations from which lab cultures were derived are highlighted in bold. C) Survival at low salinity measured 2 and 24 hours after exposure to treatment salinities, response curves are predicted from the *glmer*, colored points denote presence of active animals (0 = dead, 1 = alive), intensity denotes number of observed individuals.

## 2 RESULTS

### 2.1 Physiological results

In all populations, survival in common-garden experiments was significantly affected by treatment salinity and time (p < 0.001 for both), with higher mortality at low salinity levels and mortality increasing over time (Fig 1C, Fig. S4). Survival also differed between copepods of different origin (p < 0.001); significant interactions of origin x time (p < 0.001), and origin x treatment salinity (p = 0.015) indicated that these effects were not consistent across all conditions. When comparing model means of the low salinity treatments specifically (< 5 PSU), populations originating from low-salinity habitats (LL17, IU7, H21, and GDY) showed higher survival than other populations after 2 hours of exposure. At later timepoints, survival differences between populations from low-salinity habitats to others became less pronounced and, in some cases, reversed (e.g., H21; Fig. S4).

For the respiration experiment, model selection yielded conflicting results, where AIC favored the interaction model, but BIC supported the additive model. We therefore report results for both tests. For the simple additive model, both treatment salinity and population origin significantly affected oxygen consumption (p < 0.001 for both). However, when including an interaction term, population origin, but not treatment salinity, significantly affected oxygen consumption (p < 0.001, and p = 0.704 respectively; Fig. S5). For this model, the significant interaction between origin x salinity (p = 0.023) indicated variations in the strength and direction of the salinity effect among stations. Notably, two populations (KB03 and BB23) had significantly lower respiration rates at the low salinity treatment, while all other stations only showed non-significant decreases compared to the control.

### 2.2 Ocean connectivity and population differentiation

We assessed the oceanographic connectivity among all sampling locations using Lagrangian dispersal simulations (Fig. 2A and B). All sites, with the exception of KB03, showed a higher predicted potential for copepod retention (diagonal of Fig. 2A) than for export (off diagonal) to other regions. Within a single generation, we identified three connectivity clusters that were not interconnected with outside areas: North Sea, Western Baltic and Northern Baltic Proper (Fig. 2A). Similarly, multi-generational connectivity was highest within these connectivity clusters, while it took a maximum number of 6 simulated generations to connect all sampling locations (Fig. 2B, Fig. S6). Overall, dispersal from the North Sea towards the Eastern Baltic required more generations compared to dispersal in the opposite direction.

**Figure 2.**
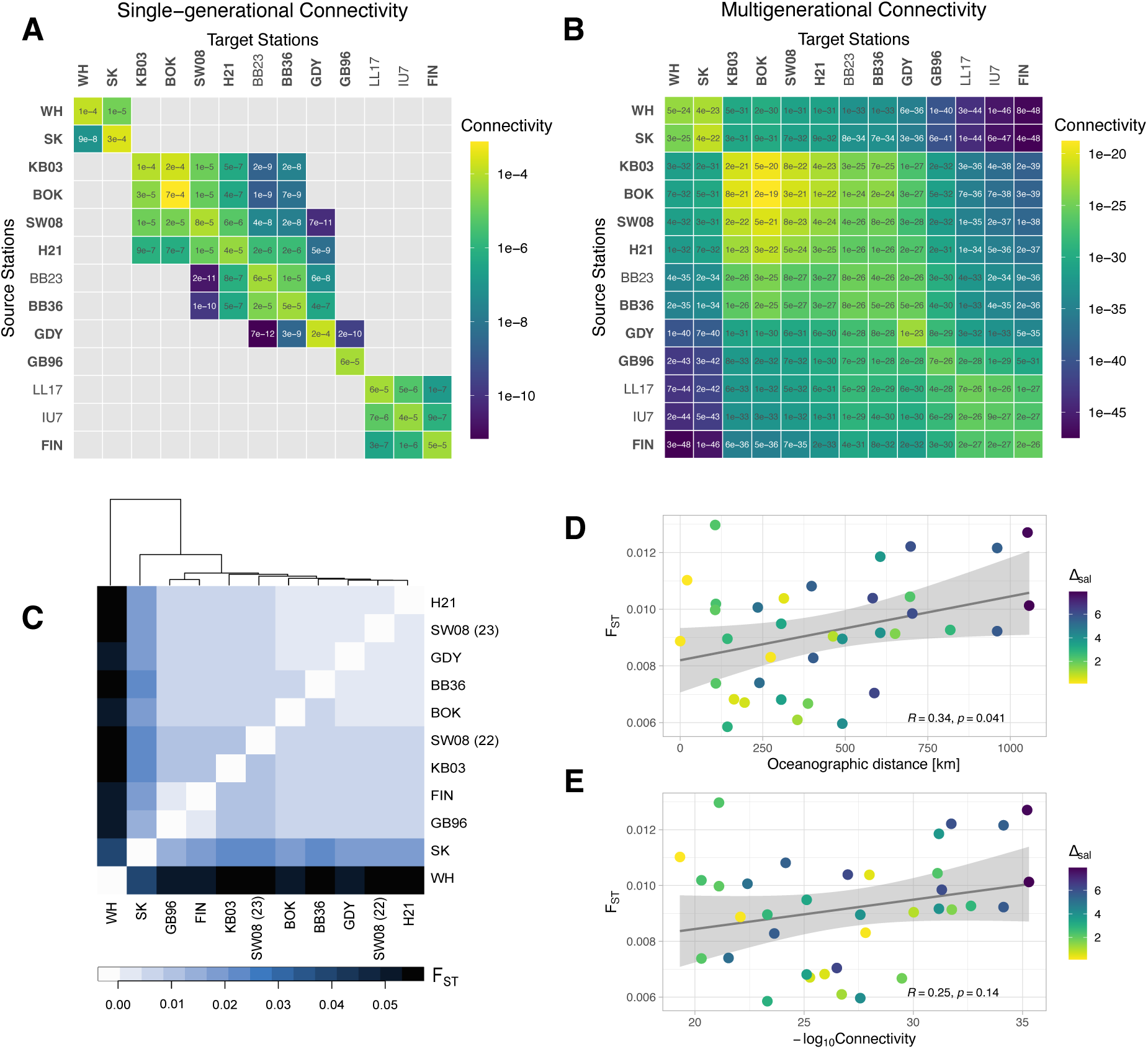
Strong population differentiation between North Sea and Baltic but weak isolation-by-distance within the Baltic. A) Single-generational connectivity between all sampling locations, connectivity is directional and measured in relative concentration of particles that are transported from a source to a target location, locations with genetic sampling are highlighted in bold. B) Multi-generational connectivity between all sampling locations C) Global pairwise F_ST_, hierarchical clustering based on similarity. E) Correlation between shortest oceanographic distance (least cost path) and F_ST_ for Baltic samples, color indicates difference in mean surface salinity between locations, results from Pearson’s correlation are shown in the plot. D) Correlation between multi-generational connectivity and F_ST_ for Baltic samples. Maximum connectivity was used for this plot, color indicates difference in mean surface salinity between locations, results from Pearson’s correlation are shown in the plot.

Estimates of F_ST_ and hierarchical clustering revealed strong genetic structure between North Sea and Baltic Sea populations consistent with oceanographic connectivity (Fig. 2C). Genome-wide pairwise F_ST_ was between 0.006 and 0.057 (mean F_ST_: 0.019 ± 0.017), with greater differentiation between North-Baltic populations (mean F_ST_: 0.038 ± 0.016) and lower differentiation within the Baltic Sea (mean F_ST_: 0.009 ± 0.002). However, there was substantial differentiation between both North Sea locations (F_ST_ = 0.041). Global nucleotide diversity was relatively high (θ_π_ = 0.016 ± 0.002) and Tajima’s D was negative for all populations (D_Taj_ =-1.90 ± 0.16), indicating an excess of rare polymorphisms (Fig. S7 and S8). There was no significant correlation of these population statistics with the salinity conditions at the sampling stations (D_Taj_: Pearson’s R = 0.051, p = 0.881; θ_π_: Pearson’s R = 0.022, p = 0.949).

Within the Baltic Sea, we found a weak but significant positive correlation of absolute distance between the sampling sites (Pearson’s R = 0.342, p = 0.041, Fig 2D), and a weak but non-significant positive correlation of multi-generational ocean connectivity with F_ST_ (Pearson’s R = 0.25, p = 0.141, Fig. 2E). Together, this suggests that isolation-by-distance is weak within the Baltic Sea. However, sampling year appeared to influence F_ST_, with higher differentiation in 2022 sample pairs than 2023 sample pairs, indicating temporal fluctuations may be significant structuring factors (Fig. S10).

### 2.3 Strong divergence between North Sea and Baltic Sea

We first identified loci potentially driving local adaptation between the North and Baltic Sea, revealing 6,202 outlier SNPs from the binary North Sea–Baltic contrast (*BayPass* BF_is_ > 20 dB, NB_contrast_). Of these outliers, 2,461 SNPs exceeded neutral expectations for genetic divergence via XtX while also pairwise F_ST_ outliers (Fig. 3A, NB_select_); these 2,461 overlapping SNPs will be referred to as putatively adaptive from this point forward.

**Figure 3.**
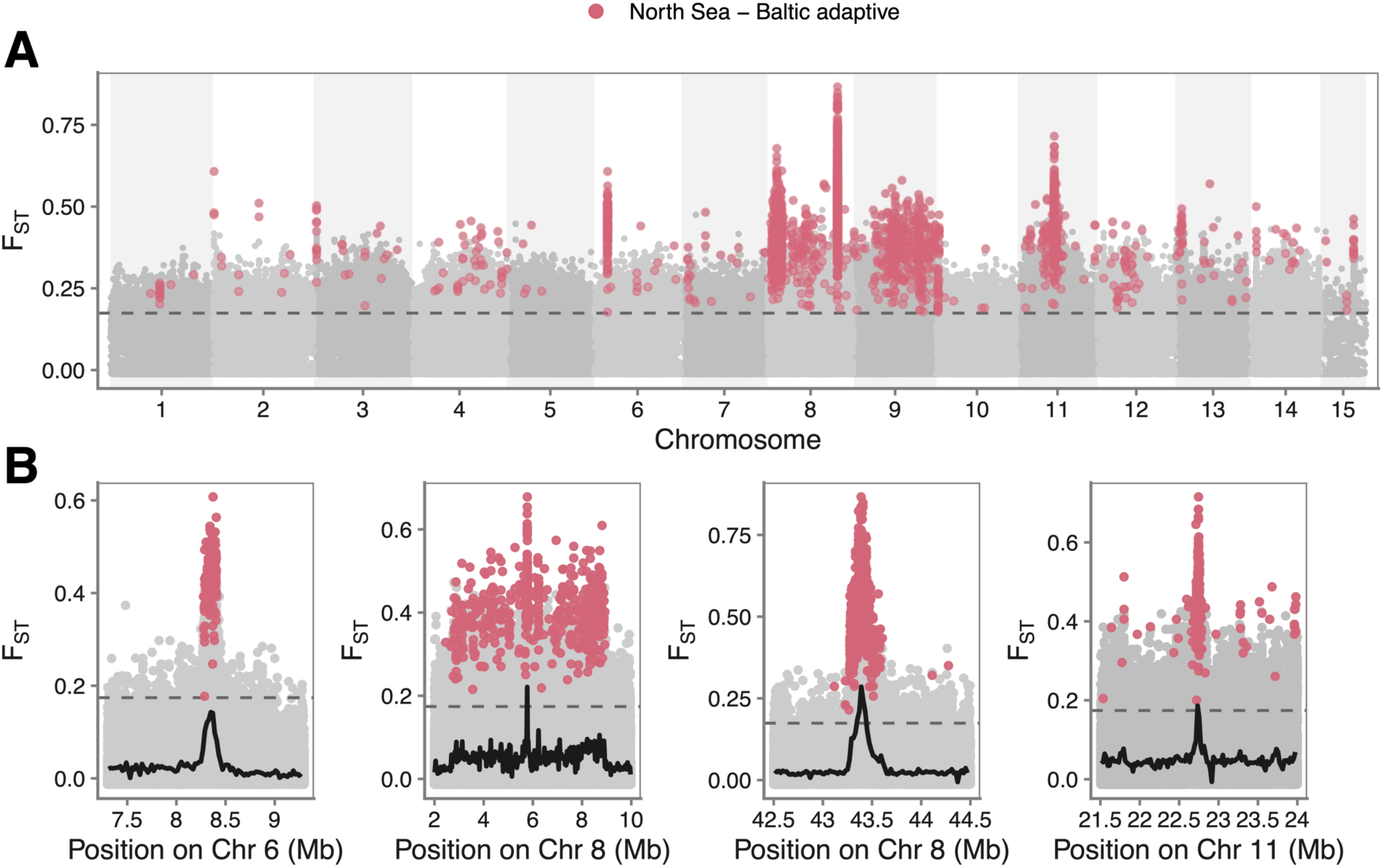
Putatively adaptive loci across North Sea and Baltic Sea. A) Manhattan plot of North Sea-Baltic per SNP pairwise F_ST_. Values above the dashed line are in the top 1 %, putatively adaptive SNPs (contrast outliers that are also under selection) are colored in magenta. B) Zoomed in view of regions of high differentiation on chromosome 6, 8, and 11, the black trend in each plot indicates F_ST_ in 20 kb.

Notably, putatively adaptive SNPs were not evenly distributed across the genome but instead strongly clustered in few regions. Out of the 15 chromosomes, four chromosomes (6, 8, 9, and 11) contained 88.5 % of all outlier SNPs (2,178 of 2,461 SNPs). Within three of these SNP-dense chromosomes, we found distinct peaks in pairwise F_ST_ (Fig. 3B). The highest peak on chromosome 6 had a maximum F_ST_ of 0.61, and was ∼150 kb in length containing 131 putatively adaptive SNPs. There were two main regions on chromosome 8, the larger one was ∼ 6.6 Mb in length and contained 520 putatively adaptive SNPs with an F_ST_ peak of 0.68, the smaller one (∼350 kb) had the highest overall pairwise F_ST_value (0.87) and contained 655 putatively adaptive SNPs. On chromosome 11, F_ST_ peaked at 0.72 within a ∼ 30 kb region that contained 134 putatively adaptive SNPs. On the other hand, chromosome 9 did not show any clear peaks and instead outlier SNPs were distributed across the majority of the chromosome (∼40 Mb). For all regions, except the chromosome 11 region, nucleotide diversity differed for most or even all populations compared to the mean q_p_ (Fig. S9). The larger region on chromosome 8 showed consistent reductions in θ_π_ in all populations, while on chromosome 6 only the Baltic populations had reduced diversity. Conversely, the small region on chromosome 8 showed reductions in nucleotide diversity for the North Sea populations (especially WH), while Baltic populations had increased θ_π_.

Overall, the putatively adaptive SNPs identified in our North Sea–Baltic contrast mapped to 600 distinct genes. In some of the regions of interest, genes could be directly linked to salinity-related processes: the 655 SNPs in the chromosome 8 peak mapped to 11 genes, involved in stress response and homeostasis. Chromosome 9 contained various key salinity genes, such as a V-type ATPase subunit 1, Na^+^/K^+^-ATPase (NKA) subunit β interacting protein, and carbonic anhydrase 2 (Table S4). Putatively adaptive genes were enriched for actin filament binding (GO:0051015) and two circulatory pathways (KEGG pathways: map04270, map04261). When taking a more lenient approach and considering all *BayPass* contrast outliers (BF_is_ > 20 dB, but no filter for F_ST_ or XtX), the 6202 SNPs mapped to 1585 genes with enrichment for G protein-coupled receptor signaling pathway (GO:0007186), as well as marginal enrichment (0.05 < adj p-value < 0.1) for calcium ion binding (GO:0005509), and protein phosphorylation (GO:0006468). In addition to circulatory pathways, this lenient set was also enriched for signal transduction pathways such as cAMP signaling (map04024) and Hippo signaling – fly (map04391, see full list in Table S5).

### 2.4 Polygenic adaptation along the Baltic Sea salinity gradient

Across the Baltic salinity gradient, adaptation was highly polygenic (Fig. 4A and B). Identifying loci overlapping between our two GEA methods, there were 4,150 mean salinity SNPs (GEA_mean_) and 6,731 salinity range (GEA_range_) SNPs, with a significant overlap of 1,605 loci between the two sets (SuperExactTest p < 0.001). The two selection scans (XtX and PCAdapt) identified 1,251 overlapping SNPs (B_select_). Allele frequencies for these outliers showed a linear correlation with the variables (Fig. 4C and D). Of these GEA outliers, 24 mean salinity outliers and 35 salinity range outliers overlapped with SNPs under selection (B_select_). Overlaps significantly exceeded chance expectations (SuperExactTest p < 0.001 for both). While this direct SNP overlap was relatively small, SNPs identified by these different methods mapped to the same genes (shared genes between B_select_ and GEA_mean_: 264 genes, shared between B_select_ and GEA_range_: 342 genes, Fig. 4 shown in yellow), indicating that different adaptive processes might be targeting different regions on the same genes.

**Figure 4.**
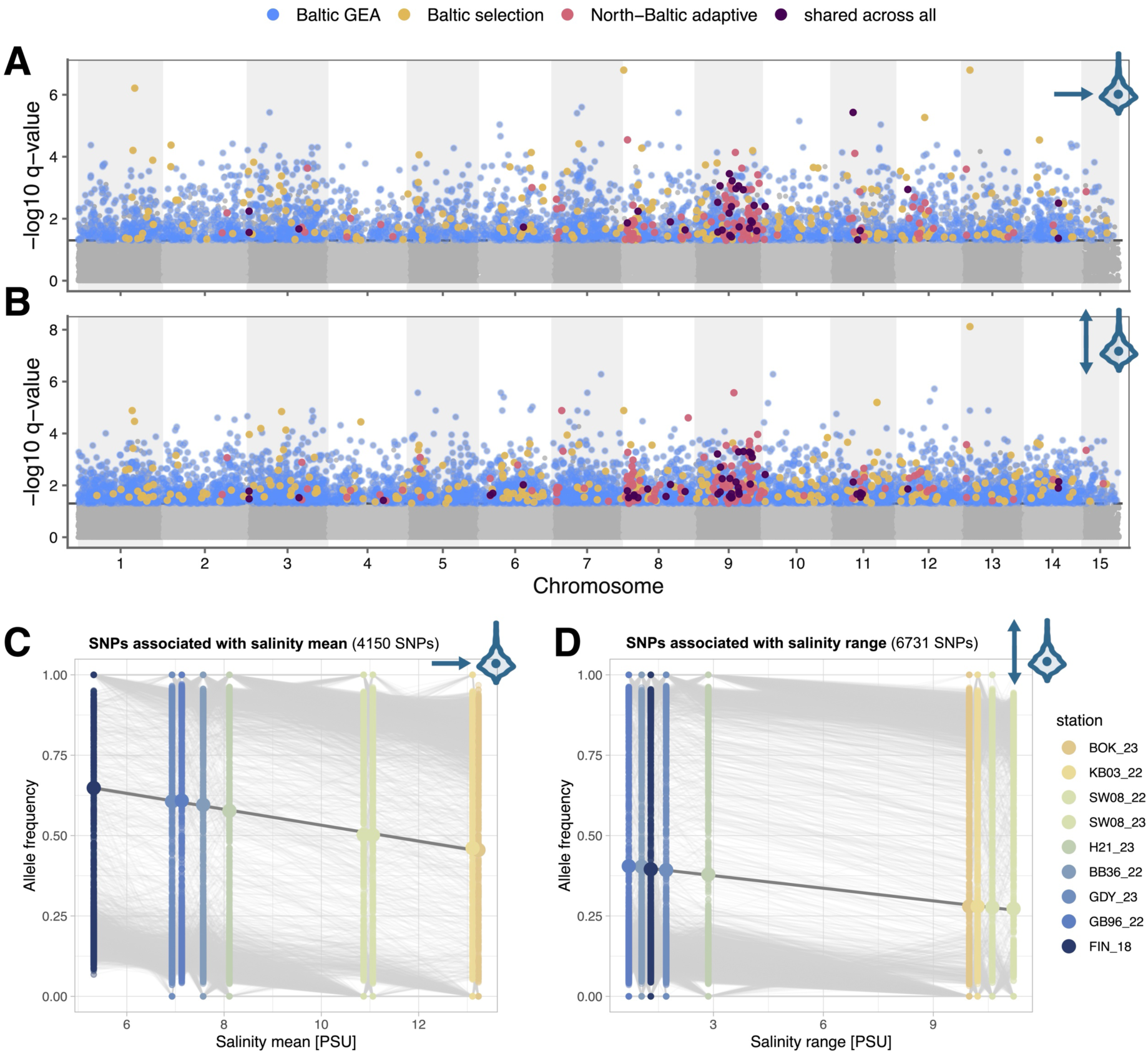
Results of genotype-environment associations for mean salinity (A, C) and salinity range (B, D). A) Manhattan plot for SNPs associated with mean salinity. Colors indicate significant GEA outliers (overlap between LFMM q-value < 0.05, and BayPass BF_mc_ > 20 dB, shown in blue), gene overlaps are highlighted by coloring the most significant SNP within the respective gene. Gene overlaps with the Baltic selection scan are shown in yellow, overlaps with North Sea – Baltic contrast analysis are shown in magenta, and genes that were found in both Baltic GEAs and all other analyses are shown in dark purple. B) Manhattan plot for SNPs associated with salinity range. Color usage is identical to A). Frequency shifts for significant GEA outliers for C) mean salinity and D) salinity range. Allele frequencies were polarized to show rising allele frequencies with decreasing salinity.

SNPs associated with mean salinity and salinity range mapped to 2,487 and 3,748 genes, respectively. For mean salinity, enriched GO terms pointed towards several functional clusters: ion and osmotic regulation (calcium ion binding, GO:0005509; voltage-gated calcium channel complex, GO:0005891, potassium ion import, GO:1990573), cell remodeling (cell-cell adhesion, GO:0098609; actin cytoskeleton organization, GO:0030036), and signaling (protein phosphorylation, GO:0006468; signal transduction, GO:0007165, protein serine/threonine kinase activity, GO:0004674). Important KEGG pathways were related to signal transduction and included cAMP (map04024), MAPK (map04010) and calcium signaling pathways (map04020). When only considering the 264 genes that were also under directional selection (shared between B_select_ and GEA_mean_), similar processes and pathways emerged with a focus on signal transduction pathways and endocrine pathways (see Table S5 for full list).

The salinity range analysis shared many of the aforementioned GO terms and pathways, but also uniquely highlighted additional enriched functions. First, transmembrane transport and ion transport processes were enriched (transmembrane transport, GO:0055085; ion transmembrane transport GO:0034220; potassium ion transmembrane transport, GO:0071805). Second, additional GO terms pointed towards neuronal signaling (G protein-coupled receptor signaling, GO:0007186; chemical synaptic transmission, GO:0007268; synapse organization, GO:0050808), suggesting a more dynamic sensory and neuroendocrine response in populations that experience fluctuating salinity conditions. KEGG pathways were mostly enriched for signaling pathways and signaling molecules. Key osmoregulatory genes that were under selection and identified as outliers by one or both GEA variables included Na^+^/K^+^-ATPase subunit β (NKA-β), K^+^-Cl^-^cotransporter (KCC), Na^+^/H^+^ exchanger (NHE), Rh protein 55, and glycine and glutamate receptors.

Unlike in the North Sea–Baltic contrast, putatively adaptive GEA SNPs within the Baltic showed no clear genomic peaks and there was little to no direct overlap between outlier loci identified by both analyses (no overlap between NB_select_ and GEA_mean_, 1 SNP between NB_select_ and GEA_range_, 2 SNPs between NB_select_ and B_select_, see Table S3). However, in many cases SNPs mapped to the same genes that were connected with the Baltic salinity gradient but also played a role in North Sea–Baltic adaptation (NB_select_ and GEA_mean_: 165 genes, NB_select_ and GEA_range:_ 204 genes, Fig. 4, shown in magenta). Finally, a small set of genes was identified across all methods: 46 genes shared between NB_select_, B_select_, and GEA_range_; 34 genes shared between NB_select_, B_select_, and GEA_mean_ (Fig. 4, shown in dark purple). Genes that were shared between the North Sea–Baltic contrast and the mean salinity GEA were enriched for protein phosphorylation and cell-cell junctions, while salinity range GEA outliers shared no significant GO terms but instead many enriched pathways with the North Sea–Baltic Sea set, among them MAPK, Ras and cAMP signaling pathways (map04010, map04014, map04024). Similarly, there was enrichment for signaling pathways for GEA variables (Calcium signaling, cAMP signaling), when only including those genes that were also shared with the Baltic selection scan and the North Sea–Baltic contrast (see Table S5 for full list).

### 2.5 Genomics offsets and habitat restrictions

Genomic offsets were most pronounced for Western Baltic populations, where the predicted salinity change was largest (Fig. 5A, see Fig. S11 for RONA), suggesting that these populations are under the greatest risk for maladaptation. Since our model approach only included mean salinity as an environmental variable, all adaptive genetic differentiation that was considered for genomic offsets was aligned with the salinity gradient. As a result, genomic offsets were reduced to the squared environmental shift from current to future salinity scaled by genomic sensitivity. However, while offsets mirror absolute change in surface salinity, relative salinity decreases might be more biologically relevant as even small absolute reductions in salinity can have a large impact in regions already approaching freshwater conditions (Fig. 5B). We therefore also developed a simple estimator of habitat suitability that derived salinity thresholds from phenotypic data (LS_5_ at 24 hours) and quantified the number of days a site exceeded these thresholds. We considered a non-adaptive and adaptive scenario for threshold determination. Under the non-adaptive scenario, Baltic *A. tonsa* would face considerable habitat loss due to salinity decline (141,436 km_2_ inviable habitat), rendering large areas of the Eastern Baltic Sea uninhabitable for the species (Fig. 5C). In contrast, the adaptive scenario showed 79 % less habitat loss (29,769 km^2^ inviable habitat) confined to the northern and easternmost distribution edges (Fig. 5D, Fig. S12).

**Figure 5.**
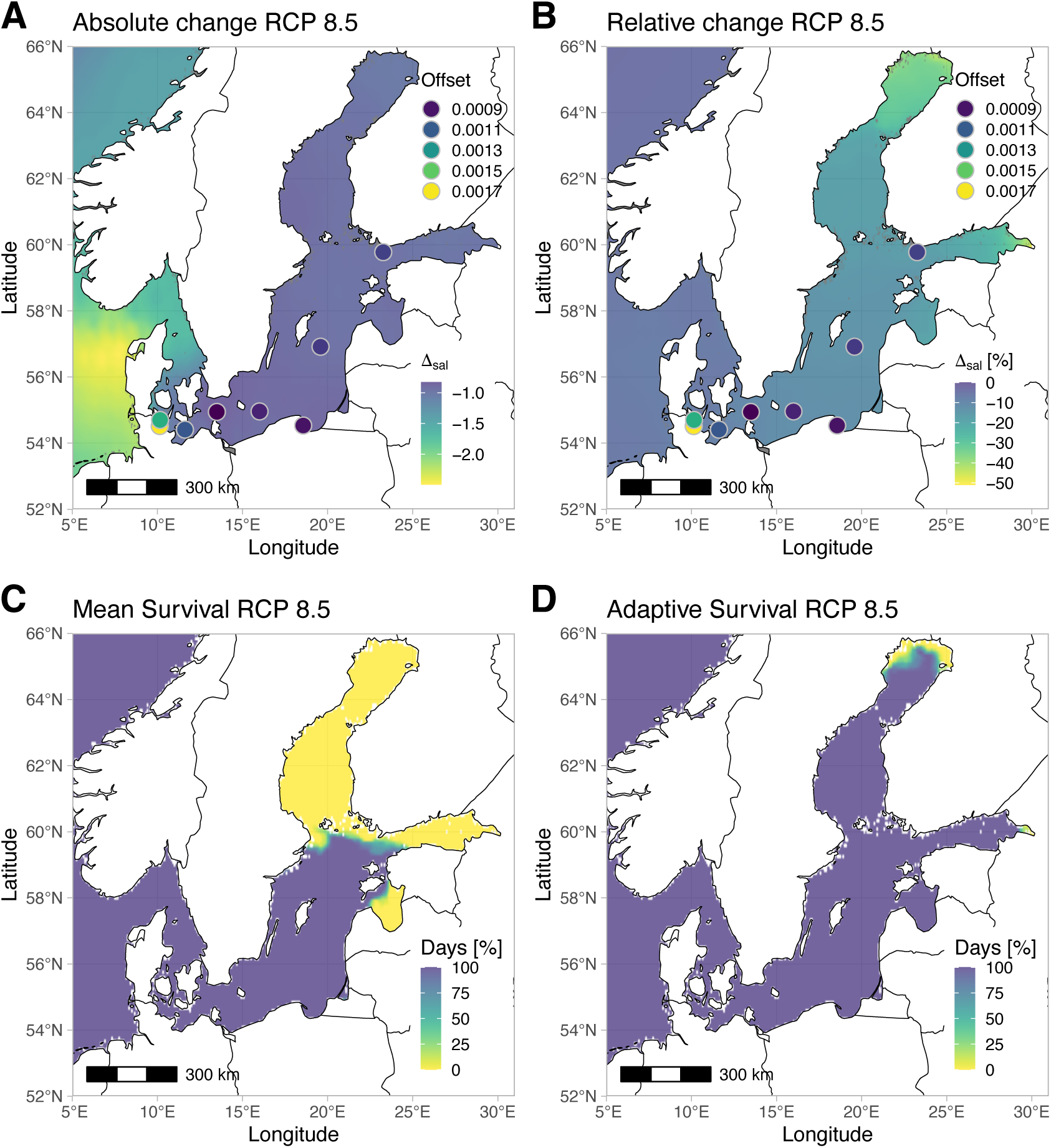
**Genomic offset and potential habitat restriction under RCP 8.5**. A) Predicted absolute salinity decline until 2100, genomic offsets per location are added as points, higher values indicate higher mismatch between current and future alleles. Notice that offsets correlate with absolute salinity change. B) Predicted relative salinity decline until 2100, with points showing genomic offset (same as in A) C) Habitat suitability expressed in percentage of days above the species’ mean salinity tolerance threshold (0% = all days below threshold, 100% = all days above threshold) as measured in our survival analysis (mean LS_5_ at 24 hours) D) Habitat suitability expressed in percentage of days above the mean tolerance threshold for the population with highest survival (LL17, LS_5_ at 24 hours). Here, we assume the greatest low-salinity tolerance represents the current adaptive maximum for the species in this region.

## 3 DISCUSSION

We combined physiological experiments, dispersal simulations, and seascape genomics to investigate how genomic architecture shapes adaptation to salinity in *Acartia tonsa* across two contrasting spatial scales: between the fully marine North Sea and the brackish Baltic Sea, and along the west-east Baltic salinity gradient. We found physiological differences between common-gardened populations originating from low and high salinity environments despite a strong dispersal potential through ocean currents, providing evidence for local differentiation in salinity tolerance. At the genomic level, differentiation between North Sea and Baltic Sea populations was primarily driven by few localized genomic regions with elevated F_ST_, whereas adaptation along the Baltic Sea salinity gradient was more broadly polygenic, with limited direct overlap with the North Sea–Baltic Sea selection signal. Together, these findings suggest that the mechanisms underlying the initial transition into the low-salinity Baltic Sea differ from those contributing to subsequent local adaptation across the Baltic salinity gradient. However, shared genes between both candidate sets were enriched for processes related to osmotic tolerance, suggesting that distinct genomic architectures may converge on common physiological pathways. Finally, by integrating physiological limits with patterns of local adaptation, we demonstrated how physiologically informed methods can improve predictions of species’ risk and resilience to changing environments relative to genomic offsets alone.

### 3.1 Physiological differences are consistent with local adaptation to salinity

Survival under low-salinity stress was consistent with local adaptation, where populations from low-salinity regions outperformed high-salinity populations. However, this was most pronounced in the short-term exposure (especially at 2 hours, but also 24 hours), while at later time points (48 and 72 hours) there was no consistent survival advantage of low-salinity populations over high-salinity populations (Fig. 1C, see also Fig. S4). This difference between responses to acute and prolonged hypo-osmotic stress might be explained by the conditions copepods face in their natural habitats: Transient salinity reductions, such as those caused by heavy rainfalls (Drushka et al., 2016; Gassen et al., 2024), represent realistic and frequently encountered challenges especially in the low-salinity regions of the Baltic, and the ability to survive these fluctuations in the instance they occur is likely under strong directional selection. The early timepoints in our experiment best captured this acute response. Conversely, prolonged continuous exposure to near-freshwater conditions for several days is less common in the typical range of *A. tonsa* as copepods can avoid experiencing sustained surface hypo-salinity by migrations to deeper saline layers (Lance, 1962). The weakening of population differences at later time-points may therefore reflect not an absence of local adaptation, but rather that prolonged continuous exposure to hypo-salinity exceeds the ecologically relevant timescale and physiological limits for the species.

In addition, we observed that in some cases, those populations that showed clear survival advantages at early time points, were among the worst performing populations later in the experiment (e.g., H21). This pattern may illustrate the high energetic cost of surviving extreme hypo-salinity. Similarly, a previous study has demonstrated that starvation for more than two days significantly decreases thermal limits in *A. tonsa* compared with non-starved individuals (Rueda Moreno & Sasaki, 2023). In our study, the observed pattern may suggest a trade-off in which copepods that allocate substantial energy reserves toward surviving low-salinity stress early may experience energetic depletion, reducing their ability to survive later in the experiment. Since we only tested salinity tolerance in the absence of food, we cannot say if low-salinity populations would be able to maintain elevated survival rates when provided with sufficient resources.

We expected that salinity reductions would correspond to an increase in respiration rates in *A. tonsa,* as has been found previously (Lance, 1965). However, our results did not align with these findings (Fig. S4). Instead, respiration rates showed slight decreases across most populations with decreasing salinity, though these patterns were only significant in two instances. Similarly, Calliari et al. (Calliari et al., 2006) found a dome-shaped respiration pattern in *A. tonsa* across a broad salinity range, with highest rates around 20 PSU and significant decreases under extreme high and extreme low salinity. The observed decrease of respiration under low-salinity stress could be explained by changes in swimming behavior. In the copepod *Eurytemora affinis*, a decrease in salinity correlated to slower swimming speeds and an increase in sinking behavior (Seuront, 2006). While we did not directly quantify swimming speeds, we observed anecdotal reductions in activity in the low-salinity treatment. Previous experiments on the swimming behavior of *A. tonsa* at low salinity did not find a reduction in movement but showed an increase in body volume that resulted in faster sinking speeds (Svetlichny & Hubareva, 2014). It is worth noting that unlike in our experiments, animals in the study by Svetlichny and Hubareva were slowly acclimated to the treatment conditions and never food limited which could mask the effects of salinity stress (Calliari et al., 2006). Future research would benefit from quantifying differences in the swimming behavior of *A. tonsa* from different salinity regimes under salinity stress in the presence and absence of food, as well as testing more drastic reductions in salinity to elicit a stronger stress response.

### 3.2 Ocean connectivity is a weak predictor of population differentiation

There was strong genetic differentiation between the North and Baltic Sea but only weak differentiation within the Baltic Sea. Our dispersal simulations confirmed the separation between Baltic and North Sea where multiple generations were necessary to connect populations. In addition, the high F_ST_ between the two North Sea populations could partly be explained by the dispersal model, where dispersal between the North Sea locations was limited due to the strong oceanographic transport boundary created by the river Elbe outflow. Within the Baltic Sea, populations were highly connected by ocean currents leading to a high potential gene flow. Most locations within the Western and Central Baltic connected within a single generation, explaining the low F_ST_ between relatively distant sites. However, connectivity was only a weak predictor of genetic differentiation. and a simple isolation-by-distance model was a better predictor of genetic differentiation, suggesting that realized gene flow might be affected by ecological factors such as salinity that are distinct from dispersal processes alone. In addition, there was an effect of sampling year on F_ST_ within the Baltic Sea. This could be attributed to the ‘egg-bank’ effect known in other diapausing species such as *Daphnia*, where annual differences in genetic composition of populations may be due to the stochastic recruitment from dormant eggs (Pfrender & Lynch, 2000; Huang et al., 2022). The contribution of sampling year to pairwise F_ST_ potentially reflects stochastic variation in egg bank cohort composition and suggests background of temporal genetic noise in our system. Regardless, the identification of consistent GEA outliers coupled with physiological evidence implies that selection along the salinity gradient is sufficiently strong to maintain local adaptation across space and time despite high gene flow and temporal variation.

The negative Tajima’s D values observed across all populations could indicate a recent population expansion leading to an excess of low frequency polymorphisms. This supports the theory that *A. tonsa* might not be native to European waters and instead was introduced to the North Sea in the early 1900s from where it spread to the adjacent Baltic Sea (Brylinski, 1981; Belmonte, 2025). Indeed, shortly after the first records in the area, the species had established itself as a major player within the zooplankton community indicating rapid population expansion (Redeke, 1934). Similarly, in the Mediterranean rapid population growth of the invasive scyphozoan *Rhopilema nomadica* within the last century has left a fingerprint of negative Tajima’s D in COI sequences (Giallongo et al., 2021). A global, phylogenetic analysis of *A. tonsa* could help to reveal native origins and order of establishment of this species; additional work is needed to resolve this question.

### 3.3 Contrasting genomic architectures reflect distance to salinity optimum

The distance of a population from its phenotypic optimum may dictate which processes dominate adaptation: large-effect loci are favoured when the required phenotypic shift is large, while populations close to their local optimum are expected to adapt through small allele frequency shifts distributed across many loci (Kopp & Hermisson, 2007; Jain & Devi, 2018; Barghi et al., 2020). The abrupt transition from fully marine North Sea conditions to the brackish Baltic represents a large phenotypic shift, while populations along the more gradual internal Baltic gradient may be closer to their local salinity optima. Accordingly, we find that the transition into the Baltic Sea is driven by few large genomic regions while adaptation within the Baltic is highly polygenic.

At the larger scale, it has been hypothesized that systems with high gene flow and strong selection should favor few and tightly linked loci instead of many small-effect loci (Yeaman & Whitlock, 2011). Chromosomal inversions are one genomic mechanism that can realize this architecture by reducing recombination and creating haplotype blocks, also known as ‘supergenes’ (Wellenreuther et al., 2025). In the North Sea–Baltic Sea transition zone inversions have been found to facilitate adaptation and their frequencies are highly correlated to the salinity gradient in herring, European plaice, and cod (Johannesson et al., 2020). Our results suggest a similar process might underlie divergence between North Sea and Baltic *A. tonsa*. We found sudden, extremely localized, increases in F_ST_ that coincided with selection scan outliers. These strongly differentiated regions contained the majority of putatively adaptive SNPs despite only representing a fraction of the genome and could be caused by structural variants such as chromosomal inversions. However, the shape of the F_ST_ peaks in our study mostly did not follow the typical inversion patterns, where inversions are expected to leave a characteristic ‘block’ pattern with highest F_ST_ values around the inversion breakpoints due to suppressed recombination within the inverted region. Instead, we often found ‘mountain’-shaped peaks that could be explained by genetic hitchhiking, where one gene is under strong selection and surrounding regions are dragged along. Similarly, Akopyan et al. (Akopyan et al., 2025) found sharp-peaked patterns in locally adapted populations of Atlantic silverside that the authors contribute to centromeric regions with reduced recombination. In the *E. affinis* species complex, ion transport genes have been found near centromeric regions caused by gene fusions (Du et al., 2025). This process where osmoregulatory genes were preserved in low recombination regions might have contributed to the species’ successful invasions of freshwater habitats and similar processes could support *A. tonsa*’s transition from marine to brackish habitats. Due to the limitations of pool-seq data, we could not definitively confirm the process underlying these peaks of divergence; additional long-read or HiC data would help resolve these processes. With regards to gene functions, multiple of the regions of interest could be linked to salinity-related processes (see Supplementary Discussion). Together with the combined evidence of strong genetic differentiation between basins, and changes in nucleotide diversity within the candidate regions, this provides strong support for a role of these regions in local adaptation.

At the smaller scale, adaptation along the internal Baltic salinity gradient followed a contrasting and highly polygenic pattern. Given the strong divergence signal and putative structural variants driving the North Sea–Baltic transition, we initially expected adaptation along the Baltic gradient to involve similar localized genomic regions, perhaps with less pronounced differentiation reflecting the smaller environmental contrast. Instead, salinity adaptation within the Baltic was highly polygenic, with GEA candidate loci distributed across the genome and none of the strongly differentiated regions from the North Sea–Baltic contrast emerging as major outliers. The signal of adaptation was clearly discernible despite the high oceanographic connectivity between the Western and Central Baltic populations and low pairwise F_ST_, which indicates high gene flow between these regions. A growing body of theoretical and empirical work (Yeaman, 2015; Barghi et al., 2020; Fuhrmann et al., 2023; Brauer et al., 2026) suggests this polygenic architecture may be common and can be maintained even in the presence of substantial gene flow. The salinity gradient in the Baltic Sea might be especially suited for this mode of adaptive architecture. Tolerating hypo-osmotic stress and large salinity fluctuations requires coordinated responses across connected physiological pathways (Boeuf & Payan, 2001). When an adaptive trait is underpinned by many genes, selection pressure is distributed across the genome instead of favoring few loci, maintaining the polygenic signal. This is consistent with our findings, where adaptation to the Baltic salinity gradient is driven by many genes contributing to different osmoregulatory processes and pathways. Similarly, polygenic adaptation has been shown in *E. affinis* adapting to the same salinity gradient (Diaz et al., 2026).

### 3.4 Common pathways and genes reveal core mechanisms of salinity adaptation

Despite the differences in genomic architectures between the North Sea–Baltic Sea contrast and the within Baltic adaptation and the unique genes associated with these architectures (see Supplemental Discussion), both comparisons also identified overlapping sets of genes suggesting shared mechanisms of adaptation. A considerable number of genes under selection in the North Sea–Baltic contrast was shared with our Baltic GEA results (165 for GEA_mean_, 204 for GEA_range_, see Table S3), an overlap that exceeded chance expectations. The observed overlap occurs almost entirely at the gene level as opposed to individual SNPs, which suggests that different populations, or selective contexts, may use distinct variants within the same adaptive genes. It is worth noting that salinity is not the only environmental factor differing between North Sea and Baltic. Other factors, such as the presence of tides may also be under selection and could therefore reduce the overlap of the selection scan with the salinity-focused GEA analyses. Regardless, the gene level convergence across methods strengthens the support for these genes as targets of salinity-related selection, and suggests that independent variants within the same genes contribute to adaptation within the Baltic and between oceanographic regions.

Across both comparisons there was enrichment for multiple pathways linked to signal transduction, including MAPK, cAMP, and Ras signaling pathways. These pathways can regulate gene expression under osmotic stress and are considered crucial in responding to changes in salinity (Dildar et al., 2025; Huang et al., 2026). In the Pacific white shrimp (*Litopenaeus vannamei*), Ras pathway genes were downregulated under low-salinity stress, likely supporting homeostasis maintenance (Farhadi et al., 2022). Similarly, chronic low-salinity stress led to enrichment in several signal transduction pathways (among them MAPK, cAMP, and Ras signaling) in a transcriptomic study on the black tiger shrimp (Li et al., 2023). Together with our findings, this strengthens the role of these signal transduction pathways in crustacean osmoregulation.

Finally, we identified Na^+^/K^+^-ATPases as likely critical genes underlying salinity adaptation. Na^+^/K^+^-ATPases (NKAs) are transporters that maintain osmotic homeostasis under fluctuating salinity (Dildar et al., 2025). In copepods, NKA has been identified as a key osmoregulatory gene in the calanoid copepod *E. affinis*: In a common-garden experiment, freshwater and saltwater inbred lines showed evolutionary shifts in gene expression of NKA (Posavi et al., 2020), and in the same species multiple subunits of NKA were under selection in parallel freshwater invasions of wild populations along the US east coast (Stern & Lee, 2020). Similarly, NKA subunits were under selection in an experimental evolution experiment testing adaptation of *E. affinis* to salinity declines in the Baltic Sea (Stern et al., 2022). NKA subunits or interacting proteins were present as outliers in all our analyses, though not all analyses consistently identified the same genes (GEA_mean_: 5 genes, GEA_range_: 6 genes, B_select_: 3 genes, NB_select_: 2 genes). This suggests that NKA is a core gene enabling low salinity adaptation across copepods and selection on NKA may have contributed to the North Sea–Baltic Sea transition and could support species persistence at low-salinity range margins.

### 3.5 Physiologically-informed models improve resilience forecasts

In our analysis, genomic offsets were largest where projected salinity decreases were most pronounced. This is likely a consequence of the modelling approaches used here: Because salinity was the only environmental driver included in the GEA, genetic offsets scaled proportionally with the projected salinity decline. As a result, the largest offset was assigned to the population that was predicted to experience the largest salinity decline, 13.2 PSU to 12.0 PSU. However, in *A. tonsa* a decline of 1 PSU at intermediate salinities is unlikely to have a significant fitness effect as this is well within the tolerance range for the species and within the range experienced by this focal population. Further, genomic offset models assume that populations are optimally adapted to their current environment and that selection pressure is uniform across populations from different environments (Ahrens et al., 2025). Therefore, it does not capture physiological thresholds or tolerance limits such as the species range margins in the northeastern Baltic Sea. In these low-salinity regions, a salinity decrease of 1 PSU could lead to physiological impairment and population collapse while the same reduction would likely have little effect in the Western Baltic. Consequently, we believe it is unlikely that our offset estimates correspond to accurate biological risks for the populations.

We therefore developed an alternative estimator of future habitat suitability at range margins that takes experimentally tested salinity limits and adaptive capacity into account. With this approach we demonstrated that even minor salinity declines could potentially lead to habitat restrictions in the northeastern Baltic Sea;large areas of this region could become unsuitable for *A. tonsa*, when considering the species’ mean salinity tolerance (non-adaptive scenario). However, when allowing for adaptation to local conditions, where we assume the populations at range margins will match the best performing low salinity tolerance in our experiments, *A. tonsa* was predicted to persist in low-salinity regions. We believe that this physiological informed predictor is a more accurate estimate of species risk as compared to genomic offsets when a threshold/range margin is involved.

It is important to note that our estimate has limitations due to three main reasons: First, because we estimate persistence based on short-term experimental data, we do not know the impact of prolonged exposure on the species fitness. In addition, we chose 5% mortality after 24 hours of exposure as a tolerance cutoff as it would signify a considerable population decline. While these thresholds accurately capture seasonal and daily weather events and significantly impact fitness (Fig. 1), we acknowledge that changing these thresholds would change the habitat suitability estimates. However, the broad finding that incorporating physiology-informed thresholds and adaptive potential into habitat suitability models improves predictions remains unchanged. Second, survival thresholds were based on adults and we currently cannot extrapolate to juveniles or larval stages. Generally, crustacean larval stages are more vulnerable compared to adults (Magouz et al., 2021; Nour et al., 2021), and under experimental conditions *A. tonsa* nauplii from the Western Baltic did not survive at a salinity of 7 PSU (Hahn et al., 2026). Therefore, even if adult individuals could survive future salinity conditions this does not mean that the reproductive cycle will be closed and the population will persist. Third, our analysis aimed to uncover adaptation to salinity and we did not include other important stressors such as temperature or hypoxia. Future studies should focus on assessing the effects of long-term exposure on all developmental stages of *A. tonsa*, as well as testing how multiple stressors interact. Previous work has shown that multiple stressors can act synergistically in *A. tonsa* (Brennan et al., 2022) and understanding the interactions between stressors such as warming and desalination would be key to predict how *A. tonsa* will react to future climatic conditions.

Previous species distribution models (SDMs) that rely on current distribution data suggest that the Baltic Sea overall will become more suitable for *A. tonsa* under climate change (Serandour et al., 2024). The authors attribute this to the increase in temperature which will benefit the warm-water species. Nonetheless, this study also shows that in coastal regions in the Central and Eastern Baltic, especially in the Gulf of Finland, habitat suitability will decline. Coincidentally, they project the largest salinity decline for the Gulf of Finland, suggesting that desalination will pose a problem for *A. tonsa.* However, databases like the Ocean Biodiversity Information System (OBIS) used to inform SDMs can introduce biases due to spatial bias in the data available (Beck et al., 2014). In the case of *A. tonsa*, data points from the (Eastern) Baltic are overrepresented in OBIS where ∼10,700 of 16,870 records are within the Baltic, despite the species’ global distribution, which may lead to skewed estimations of the species’ salinity tolerance. Our survival experiments demonstrate that while low-salinity populations can tolerate lower salinities, salinity functions as a threshold trait, with drastic survival decreases even in locally adapted populations. Further, while genomic offsets by default include estimates of local adaptation, classical SDMs often do not. However, accounting for adaptation can greatly influence SDM outcomes (Bush et al., 2016). Similarly, we were able to show the strong discrepancies in habitat suitability when using non-adaptive vs. adaptive thresholds. Therefore, experimentally validated thresholds are crucial to improve SDM accuracy (Cong et al., 2025) and we advocate for the inclusion of tolerance thresholds and knowledge of local adaptation in SDMs to better assess climate effects on habitat suitability.

Finally, while multi-generational common-garden experiments allowed us to estimate tolerance thresholds, this is only possible in easily culturable organisms such as *A. tonsa.* In other systems, genomic offsets provide a valuable tool to predict species resilience and (mal)adaptation to future conditions directly from genomic data in the face of climate change (Gain et al., 2023). Further, genomic offsets allow one to test multiple environmental stressors simultaneously, which can be challenging to execute and interpret in experimental systems. In complex environments or if the main stressor is not known, this multi-stress framework can be crucial to gain a holistic understanding of population resilience (Dauphin et al., 2023).

Our results demonstrate that *A. tonsa* local adaptation to salinity occurs across multiple spatial scales and is underpinned by distinct evolutionary mechanisms and genomic architectures that allow for local adaptation despite high gene flow. The transition between the high salinity North Sea and the brackish Baltic Sea is facilitated by few, strongly differentiated genomic regions. This pattern could be explained by structural variants combined with selection acting on a small number of large-effect loci or spatially clustered loci along a sharp environmental and demographic boundary. By contrast, adaptation along the salinity gradient within the Baltic Sea follows a highly polygenic pattern, with salinity-associated variation distributed broadly across the genome. Dispersal simulations predicted high connectivity and therefore potential gene flow within the Baltic; the presence of local adaptation indicates that selection can outweigh the homogenizing effect of gene flow. Together, these results demonstrate that the genomic architectures underlying salinity adaptation are highly complex and shaped by differing selection pressures and levels of gene flow acting on the North Sea and Baltic Sea populations versus within the Baltic Sea. Our findings also highlight limitations of genomic offset approaches in systems where adaptation is governed by threshold-like environmental responses. This suggests that forecasting frameworks should incorporate experimentally validated physiological thresholds alongside information on adaptive potential and dispersal. Such integrative approaches could offer a more accurate and biologically grounded basis for predicting species’ responses to environmental change. Together, our study highlights the value of integrating approaches to improve our understanding of adaptation and strengthen forecasts of species’ resilience to future environmental change.

## 4 MATERIALS AND METHODS

### 4.1 Sampling and culturing

Samples for population genomics were collected during late summer in 2018 and 2022-2023 using 100 μm WP2 net hauls in the North and Baltic Sea capturing a wide range of salinity conditions (see Fig. 1A and B; Table S1 for overview). Animals were sorted and stored in 95% ethanol at-20 °C until processing (N = 100). From ten of the sampling locations, we simultaneously collected live animals to start laboratory cultures. All cultures were established with ∼200 adult *A. tonsa* individuals and common-gardened at 15 PSU and 15.5 °C (± 0.3°C) in a 12:12 light:dark regime. For feeding, the microalgae *Rhodomonas spp*. was grown at identical conditions and copepod cultures were fed every other day at concentrations of 500 μgC/L. Water exchanges were performed every 2 to 4 weeks using 0.2 μm filtered sea water and salinity was monitored and adjusted with a mix of deionized water and tap water. Copepods were reared at common-garden conditions for at least three generations before the experiments and all experiments were carried out at the rearing temperature. For simplification, animals originating from the same location will be referred to as ‘populations’ throughout this manuscript even though this might not reflect the true demography. For each location we calculated surface mean salinity and salinity range using the Copernicus Global Physics Ocean Reanalysis data set (https://doi.org/10.48670/moi-00021). We used daily sea surface salinity measurements for the sampling year plus five years prior to capture interannual variability (Table S1), and only considered the months from June to October, which represents a conservative estimate of the reproductive window of the species (Witalis et al., 2024; Dutz et al., 2025).

### 4.2 Physiological experiments

For the common-gardened populations we measured two physiological traits that may be indicative of local adaptation: survival and respiration. Survival and respiration rates for populations WH and SW08 have been included in a previous study (Hahn et al., 2026). Data analysis was done in R version 4.2.2 (Team, 2022).

To estimate survival under extreme low salinity conditions (hyposalinity), we tested 5 salinities plus the control salinity (0.1, 1, 2, 3, 5, and 15 PSU) by diluting filtered sea water with a mix of tap and deionized water. For each population, 90 recently matured females were acutely transferred into experimental vials (six salinities) filled with 9 mL of water, with three replicates per treatment each containing 5 individuals. Survival was monitored after 2, 24, 48 and 72 hours by creating a gentle water flow using a pipette. If animals showed no activity, they were scored as dead and if mortality at the control salinity exceeded more than one individual within the first 24 hours, the trial was repeated. For the statistical analysis, we fit a generalized linear mixed-effect model with binomial distribution using the *lme4* package (Bates et al., 2015). We used the log_10_ transformation of salinity as a continuous predictor, as the treatment salinities span several orders of magnitude. Replicate groups were included as random factors to account for repeated observations over time. Significance of fixed effects was assessed using type-III Wald tests. Predicted survival probabilities and pairwise station contrasts were obtained from estimated marginal means using the *emmeans* package (Lenth, 2023). We derived LS_50_ and LS_5_ values from the model intercepts and slopes.

To determine base respiration rates, mature females were picked and acclimated to the experimental salinities for 16 hours without food. We measured respiration rates at low salinity (5 PSU) and the culture salinity (15 PSU). On the experimental day, animals were placed in pairs into 80 μL wells at the experimental conditions (8 replicates of female pairs), after gently rinsing them with filtered seawater (0.2 μm filter). To ensure measurement consistency, two populations were run at the same time if possible and animals were randomly assigned to one of 24 wells (16 wells with animals, 8 empty controls). Oxygen concentrations were measured for up to 2 hours with a PreSens optical sensor using the SDR_v4.0.0 software. Animals were preserved in 1% Lugol’s solution to later be photographed using a Nikon imaging microscope with NIS-Elements software (v. 5.20.000). All pictures were taken with the same magnification to ensure consistency. Prosome length was then measured using the software ImageJ (Schneider et al., 2012). Respiration rates were calculated using the *respR* package (Harianto et al., 2019) and standardized per dry body weight using the length obtained from the images and the conversion factors as defined by Kiørboe et al. (Kiørboe et al., 1985). For statistical testing, we compared an additive linear model (oxygen ∼ treatment salinity + origin) to an interaction model (oxygen ∼ treatment salinity * origin) using AIC and BIC.

### 4.3 Connectivity analysis

To assess population connectivity of *A. tonsa* by ocean currents in the Baltic Sea, species-specific traits were used to parametrize Lagrangian dispersal simulations (see Supplemental Methods for full details). We used the operational circulation product HIROMB-BOOS-Model (Brüning et al., 2021). The model is forced by the operational atmospheric model of the German Weather Service (DWD), the BSH North East Atlantic model at the boundary (Brüning et al., 2014), 19 tidal constituents and river discharge forecasts (PegelOnline and EHYPE). The model covers the entire Baltic and North Sea region with two nested grids: 1) A fine grid with a ∼900 m horizontal resolution (25 vertical layers) covering the German Bight and Western Baltic Sea, and 2) a coarse grid with a horizontal resolution of ∼5 km (36 vertical layers). Data is available since 2016 in 15-minute intervals for the three-dimensional current and hourly resolution for temperature and salinity, enabling the representation of tidal dynamics and short-term environmental variability throughout the region.

The dispersal of copepods was simulated using the Lagrangian particle spreading toolbox OceanParcels (Delandmeter & van Sebille, 2019). Particles were released with spatially uniform random distribution within 5×5 km^2^ areas from surface to 25 m at the sampling locations. Two coastal sampling sites (WH and SK) were slightly shifted offshore because their original positions were too sheltered on the model grid and prevented realistic dispersal trajectories. In addition, we released particles from 15 additional, equally spaced sites in between the sampling sites in order to allow for stepping stones and the calculation of a multi-generational connectivity matrix. One thousand particles were released per site every day at 12:00 am from the 1^st^ of June to 31^st^ of October 2016-2024 to represent the reproductive season and to capture interannual variability (46,368,000 particles in total). Trajectories were recorded for 28 days in 15 min time steps. Copepods were simulated as passively drifting particles without active vertical movement since vertical swimming capacities are not well understood for the studied species. We restricted the dispersal to depths less than 25 m and re-positioned particles to 24 m if they dispersed below this point.

To quantify direct connections among locations, Voronoi polygons were constructed around all release sites including the 15 intermediate stepping stone sites (see Fig. S1). We considered sites connected if particles from a source site reached a target site’s Voronoi polygon within the reproductive window of the total lifespan, i.e., between 10 and 28 days. The number of particle positions falling inside each target polygon was normalized by (i) the water-filled polygon area to obtain particle density per 25 km², and (ii) the total number of particle-timesteps and number of released particles to derive relative concentrations per 25 km².

Relative dispersal from each source to each target site was expressed as:

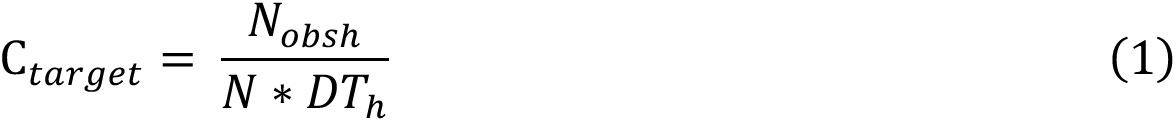

where C_target_ denotes the relative concentration (0–1) at the target site, N_obsh_ the cumulative particle-timesteps within that site, N the number of released particles, and DT_h_ the dispersal duration in 15-minute intervals (184 release days x 24 h x 4). Connectivity was averaged across all timepoints to obtain average dispersal across a diverse range of seasonal and interannual variability in large-scale weather conditions.

To explore multi-generational connectivity, single-generational connectivity matrices were iteratively multiplied and the minimal generation time linking source and target sites determined. This approach identifies dispersal corridors that may facilitate the spread of copepods over successive generations. In addition, we calculated the shortest oceanographic path and distance between all sampling stations as a simplified measure of connectivity using the *lc.dist()* function of *marmap* (Pante et al., 2023).

### 4.4 DNA extraction, sequencing and sample processing

DNA for whole genome sequencing was extracted from pooled animals (N = 100, except FIN where N = 80). Pooled sequencing provides accurate estimates of genome-wide allele frequencies, providing a cost-effective way of studying local adaptation (Futschik & Schlötterer, 2010). Animals were rehydrated in MilliQ water before homogenizing the samples with a pestle. For sample purification we used Qiagen DNeasy spin columns adding an extra wash step and RNase A treatment. Quality was assessed using Qubit and Nanodrop and samples were stored at-20 °C until sequencing. Library preparation and sequencing was conducted by the Competence Centre for Genomic Analysis Kiel (CCGA) across two lanes of an Illumina NovaSeq 6000.

*Fastp* v.0.23.4 was used for adapter trimming (Chen et al., 2018) and sample quality was checked using *FastQC*v.0.12.1 and *MultiQC* v.1.21 (Andrews, 2010; Ewels et al., 2016). We aligned to the *Acartia tonsa* reference genome (NCBI accession GCA_053477335.1) with *bwa-mem2* v.2.2.1 (Vasimuddin et al., 2019). Mapping rates and alignment quality were checked using *SAMtools* v.1.21 and *qualimap* v.2.3 (Li et al., 2009; Okonechnikov et al., 2015) and one sample was excluded after not meeting our coverage requirements (BB23). For all remaining samples, mean mapping rate was 92.19 ± 3.81% (see Table S2). Samples were deduplicated and overlaps clipped using the *dedup* and *clipOverlap* functions from *bamUtil* v.1.0.15 (Jun et al., 2015). We then identified indels using GATK v.3.8 *RealignerTargetCreater* and realigned around indels with GATKs *IndelRealigne*r. After realignment, mean sequencing depth was 78.04 ± 12.06 x. We used *grenedalf* v.0.6.3 (Czech et al., 2024) to convert the BAM files into a sync format for downstream analysis, applying a filter for alignment and base quality (Q < 30). For each population a minimum of 40 and a maximum of 160 reads were required, and a minimum of four reads were required for a base call. Diversity metrics were computed for the whole genome and in 50kb windows using *grenedalf*. For detecting SNPs, we applied a global minor allele frequency filter (MAF) > 0.05 and only retained bi-allelic SNPs. From all sync files, we excluded unplaced scaffolds, only considering the 15 chromosomes for further analysis. After applying our SNP filters, we retained 21,440,970 SNPs across all populations (North Sea and Baltic Sea), and 22,744,641 SNPs within the Baltic samples. Pairwise Hudson’s F_ST_ was computed on the filtered SNP set including all samples.

### 4.5 Detecting signals of adaptation between North Sea and Baltic Sea

We hypothesized that the transition from the high-salinity North Sea to the low-salinity Baltic Sea would involve local adaptation between these regions. To identify these loci, we combined multiple complementary approaches: First, we identified North Sea–Baltic contrast outliers using the standard covariance model with Importance Sampling (IS) in *BayPass* v.3.1 (Coop et al., 2010; Gautier, 2015) treating North Sea and Baltic Sea regions as a binary trait. To reduce computational runtime, we followed the thinning approach recommended in the *BayPass* manual, dividing the full SNP set into 99 subsets equally distributed across the genome. Each subset was run independently three times with a different random seed using 10 pilot runs of 500 iterations, a burn-in of 1000 iterations, and 5000 iterations of MCMC sampling run post burn-in sampled every 20 iterations. Convergence was confirmed by checking MCMC acceptance rates against *BayPass* target values and by verifying consistency across model runs. For each SNP, we took the median across the three independent runs as the final parameter estimate, and results from all 99 subsets were merged prior to downstream analysis. Second, we identified signals of adaptive divergence in *BayPass* utilizing the XtX differentiation statistic, which can be understood as a covariance corrected per-SNP F_ST_ (Gautier, 2015). We empirically determined significance thresholds for calibrated XtX using pseudo-observation dataset (POD) simulations, with the 99.5^th^ percentile of the POD-derived null distribution as the outlier threshold. Loci showing strong evidence of association with the North Sea–Baltic contrast were identified using a Bayes Factor threshold of BF_is_ > 20 dB, following Jeffreys (Jeffreys, 1961). Lastly, to complement the *BayPass* analysis, we calculated pairwise F_ST_ per SNP between North Sea and Baltic populations. We constructed two pooled pseudo-populations by averaging allele frequencies across all North Sea and all Baltic sampling stations respectively, converting averaged frequencies back to read counts using a depth factor of 160 (equal to the maximum depth filter) to generate a two-sample sync file. Per-SNP F_ST_ was then calculated using the ANOVA method implemented in *poolfstat (Gautier et al., 2022)*, with pseudo-population sizes set to 160 haploid individuals.

SNPs were classified as contrast outliers if they exceeded the BF_is_ > 20 dB threshold. Contrast outliers that also showed differentiation with XtX above the POD-derived significance threshold and falling within the top 1% of pairwise FST values, were considered putatively adaptive (see Fig. S2).

### 4.6 Genotype-environment association (GEA)

For the genotype-environment association (GEA), we restricted the analysis to the Baltic Sea populations to avoid masking subtle environmental associations with the strong North Sea–Baltic differentiation signal. We tested for correlations with two environmental variables: mean salinity and salinity range. Due to the significant correlation between both environmental variables (Pearson’s R = 0.569, p = 0.034), a separate analyses was run for each parameter.

We used the *BayPass* auxiliary model following the same approach as described in the previous section. To correct for population structure, the auxiliary model uses a scaled population covariance matrix which was estimated by running the base model on a randomly selected 5% of SNPs drawn from windows distributed across the genome. Bayes Factor and calibrated XtX statistics were summarized as before, with POD-derived 99.5^th^ percentile thresholds applied to the XtX statistic. In parallel, we used Latent Factor Mixed Models (LFMMs) implemented in the *LEA* package (Gain & Francois, 2021). Since LFMM cannot handle pooled samples directly, we simulated 50 haploid individuals per population from pooled allele frequencies using *rbeta* as suggested by the package developers. As with *BayPass*, analyses were run on the same 99 subsets to reduce runtime. We used the exact least-squares approach with K = 4 based on the PCA screeplot of genome-wide allele frequencies (see Fig. S3). Again, two separate models were run for the two environmental variables. Resulting p-values were calibrated using the genomic inflation factor and results from all subsets were merged prior to applying multiple testing correction using the *qvalue* package with a false discovery rate (FDR) < 0.05 (Storey et al., 2022). Finally, to detect selection independently from environmental variables, we used *PCAdapt* (Luu et al., 2017) in pool-seq mode with K = 4, again applying q-value correction with FDR < 0.05.

Loci were classified as GEA outliers if they showed strong evidence of environmental association in both *BayPass*(BF_mc_ > 20 dB) and LFMM (FDR < 0.05). Loci were classified as selection outliers if they exceeded the POD-derived XtX threshold and were significant *PCAdapt* outliers (FDR < 0.05). SNPs meeting both of these criteria (GEA outliers and selection outliers) were considered putatively adaptive. The statistical significance of overlaps between methods was assessed using the *SuperExactTest* package in R.

### 4.7 Gene mapping and functional enrichment

To investigate gene identity and functional enrichment, candidate SNPs were linked to the NCBI RefSeq annotation (GCF_053477335.1) using *BEDTools closest (Quinlan & Hall, 2010)*. Only genes within 5kb up-or downstream of a SNP were considered for functional enrichment. Gene names reported in the text follow NCBI nomenclature. Gene ontology enrichment (GO) was performed with the *enricher* function of *clusterProfiler (Wu et al., 2021)* and a gene background constructed from all SNPs present in our analysis. Results were visualized using the *dotplot* function. Pathway enrichment based on the Kyoto Encyclopedia of Genes and Genomes (KEGG) was performed with the enricher function using a custom annotation built with EggNOG-mapper v. 2.1.13 (Cantalapiedra et al., 2021).

For the North Sea–Baltic contrast analysis, we tested enrichment for the putatively adaptive set as well as the contrast outliers (BF_is_ > 20 dB). Within the Baltic analysis, we performed GO and KEGG enrichment on the GEA outliers for mean salinity and salinity range, and on genes that were identified by selection scans. There was little direct overlap of SNPs identified by several independent methods, however in many cases outlier SNPs from different analyses mapped to the same genes. Therefore, we also investigated GO enrichment of genes shared between two or more analyses. To assess the functional enrichment of genes that were driving salinity adaptation and were under directional selection, we used the gene overlap between GEA and selection scan outliers. Since we expected that adaptive processes between North Sea and Baltic, i.e., adaptations that facilitate inversion into brackish waters could be the same processes that allow persistence across the Baltic salinity gradient, we assessed the overlap between North Sea–Baltic contrast outliers and Baltic GEA outliers. Finally, we manually screened outlier genes for genes related to salinity adaptation and osmoregulation in the copepod literature (Posavi et al., 2014; Lee, 2021; Du et al., 2025).

### 4.8. Genomic offset and viability modelling

We calculated genomic offsets and RONA (risk of non-adaptedness) using *compute_genetic_offset()* function in *BayPass*. While it appears that there is local adaptation across space for *A. tonsa*, subtle shifts in salinity in the core of the salinity range are physiologically unlikely to result in strong fitness reductions for this species given their high plasticity and gene flow (see Fig. 1 and 2). Further, genomic offsets perform suboptimal for populations close to their range margins (Ahrens et al., 2025). Therefore, we developed an approach to estimate changes in habitat suitability under future salinity declines based on experimentally determined physiological limits, incorporating our insights on adaptive potential. To do this, we used our physiological data to determine the salinity at which 5 % mortality will occur within 24 hours (LS_5_) with the reasoning that this mortality rate would constitute a dramatic fitness reduction and 24 hours is an ecologically relevant timescale given local weather events and accounting for the species’ diel vertical migrations. We determined LS_5_ under a non-adaptive scenario (using the predicted mean for all tested populations) and the adaptive scenario (using the predicted mean for the best performing population, LL17) where we assumed the best performing population likely represents the range limit and maximum low-salinity tolerance within this region. Since most available climate models lack fine-scale temporal resolution, we joined contemporary salinity data from Copernicus with the BioOracle3 RCP 8.5 forecast for 2100 by nearest coordinate to estimate daily future sea surface salinity. For each coordinate point, we then calculated the percentage of days above the LS_5_ value as a proxy for habitat suitability. We used the area for those grid cells that had 0 % habitat suitability as an absolute measurement of inviable habitat.

## Acknowledgments

We thank Diana Gill, Fabian Wendt, and Claas Hiebenthal for their technical support in the lab and climate chambers, and Frank Melzner, Thorsten Reusch, and Meike Stumpp for their scientific counsel. We thank Pierre De Wit for his input on the manuscript. We are grateful for the crew and scientists aboard the RV Alkor cruises AL580 and AL601 (chief scientist: Felix Mittermayer-Schmittmann) and RV Aranda cruise COMBINE3 (chief scientist: Pekka Kotilainen); warm thanks to Siru Tasala for her help with sampling. We are grateful for Jasmin Renz and Astra Labuce for providing additional preserved animal samples (even though we could not include the Riga sample). This research was supported in part through high-performance computing resources available at the Kiel University Computing Centre. Sequencing was done in cooperation with the Competence Centre for Genome Analysis Kiel (CCGA), we thank Sören Franzenburg for the good communication. Large language models (LMMs) have been used for assistance with coding and copyediting; all generated outputs have been verified. The scientific results and conclusions, as well as any views or opinions expressed herein, are those of the authors and do not necessarily reflect those of NOAA or the Department of Commerce.

## Author contributions

AH and RSB designed this study. AH, and RSB, collected the animals. GA, JCNS and AH cultured animals. AH conducted the experiments with assistance from JCNS, GA and RSB. AH processed, analyzed, and visualized the data with input from RSB. JR, LS, and WR carried out the dispersal analysis. AH wrote the manuscript with contributions from RSB. All authors edited and approved the final version of the manuscript.

## Data, code, and materials availability

The code and physiological data that support the findings of this study are archived and openly available on Zenodo (https://doi.org/10.5281/zenodo.21358410). Whole genome sequences and sample metadata is available at NCBI under the project accession PRJNA1314997.

Hydrodynamic model data is publicly available via the Bundesamt für Seeschifffahrt und Hydrographie (BSH, https://gdi.bsh.de/de/feed/Data-of-the-Operational-Models.xml) under the open access license (https://www.govdata.de/dl-de/by-2-0). Notebooks to reproduce Lagrangian dispersal simulations are available on Zenodo (https://doi.org/10.5281/zenodo.18389990). Particle trajectories are not provided due to their large size (>4TB) but can be made available upon request.

## REFERENCES

Ahrens, C. W., Rymer, P. D., & Miller, A. D. (2025). Genetic offset and vulnerability modelling under climate change scenarios: common misinterpretations and violations of evolutionary principles. Evolution. 10.1093/evolut/qpaf216

Akopyan, M., Tigano, A., Jacobs, A., Wilder, A. P., & Therkildsen, N. O. (2025). Genetic Differentiation is Constrained to Chromosomal Inversions and Putative Centromeres in Locally Adapted Populations With Higher Gene Flow. Molecular Biology and Evolution, 42(5), msaf092. 10.1093/molbev/msaf092

Andrews, S. (2010). FastQC: a quality control tool for high throughput sequence data. In: Cambridge, United Kingdom.

Barghi, N., Hermisson, J., & Schlötterer, C. (2020). Polygenic adaptation: a unifying framework to understand positive selection. Nature Reviews Genetics, 21(12), 769–781. 10.1038/s41576-020-0250-z

Bates, D., Mächler, M., Bolker, B., & Walker, S. (2015). Fitting Linear Mixed-Effects Models Using lme4. Journal of Statistical Software, 67(1), 1–48. 10.18637/jss.v067.i01

Beck, J., Böller, M., Erhardt, A., & Schwanghart, W. (2014). Spatial bias in the GBIF database and its effect on modeling species’ geographic distributions. Ecological Informatics, 19, 10–15. 10.1016/j.ecoinf.2013.11.002

Belmonte, G. (2025). Pelagic Lifestyles in Confined Environments. In G. Belmonte (Ed.), Elements of Pelagos Biology (pp. 173–216). Springer Nature Switzerland. 10.1007/978-3-031-74280-4_5

Boeuf, G., & Payan, P. (2001). How should salinity influence fish growth? Comp Biochem Physiol C Toxicol Pharmacol, 130(4), 411–423. 10.1016/s1532-0456(01)00268-x

Brauer, C. J., Bertram, A., Sandoval-Castillo, J., Fowler, A., Bell, J. D., Hamer, P., Wellenreuther, M., & Beheregaray, L. B. (2026). Environmental Gradients Decouple Demographic and Adaptive Connectivity in a Highly Mobile Coastal Marine Species. Mol Ecol, 35(5), e70273. 10.1111/mec.70273

Brennan, R. S., deMayo, J. A., Dam, H. G., Finiguerra, M., Baumann, H., Buffalo, V., & Pespeni, M. H. (2022). Experimental evolution reveals the synergistic genomic mechanisms of adaptation to ocean warming and acidification in a marine copepod. Proceedings of the National Academy of Sciences of the United States of America, 119(38), e2201521119. 10.1073/pnas.2201521119

Brennan, R. S., Healy, T. M., Bryant, H. J., La, M. V., Schulte, P. M., & Whitehead, A. (2018). Integrative Population and Physiological Genomics Reveals Mechanisms of Adaptation in Killifish. Mol Biol Evol, 35(11), 2639–2653. 10.1093/molbev/msy154

Brüning, T., Janssen, F., Kleine, E., Komo, H., Foerderreuther, S., Menzenhauer-Schumacher, I., Jt, S., & Dick, S. (2014). Operational ocean forecasting for German coastal waters. Die Küste, 81, 273–290.

Brüning, T., Li, X., Schwichtenberg, F., & Lorkowski, I. (2021). An operational, assimilative model system for hydrodynamic and biogeochemical applications for German coastal waters. Journal of Applied Hydrography, 6-15. 10.23784/HN118-01

Brylinski, J. M. (1981). Report on the presence of *Acartia tonsa* Dana (*Copepoda*) in the harbour of Dunkirk (France) and its geographical distribution in Europe. Journal of Plankton Research, 3(2), 255–260. 10.1093/plankt/3.2.255

Bush, A., Mokany, K., Catullo, R., Hoffmann, A., Kellermann, V., Sgro, C., McEvey, S., & Ferrier, S. (2016). Incorporating evolutionary adaptation in species distribution modelling reduces projected vulnerability to climate change. Ecol Lett, 19(12), 1468–1478. 10.1111/ele.12696

Calliari, D., Andersen, C. M., Thor, P., Gorokhova, E., & Tiselius, P. (2006). Salinity modulates the energy balance and reproductive success of co-occurring copepods Acartia tonsa and A. clausi in different ways. Marine Ecology Progress Series, 312, 177–188. 10.3354/meps312177

Cantalapiedra, C. P., Hernandez-Plaza, A., Letunic, I., Bork, P., & Huerta-Cepas, J. (2021). eggNOG-mapper v2: Functional Annotation, Orthology Assignments, and Domain Prediction at the Metagenomic Scale. Mol Biol Evol, 38(12), 5825–5829. 10.1093/molbev/msab293

Cervetto, G., Gaudy, R., & Pagano, M. (1999). Influence of salinity on the distribution of *Acartia tonsa* (*Copepoda*, Calanoida). Journal of Experimental Marine Biology and Ecology, 239(1), 33–45. 10.1016/s0022-0981(99)00023-4

Chen, S., Zhou, Y., Chen, Y., & Gu, J. (2018). fastp: an ultra-fast all-in-one FASTQ preprocessor. Bioinformatics, 34(17), i884–i890. 10.1093/bioinformatics/bty560

Cong, J., Tao, Z., Xu, Y., & Li, X. (2025). From thermal tolerance to distribution resilience: Physiology-informed species distribution models moderated range shifts for an intertidal crab species under climate change. Global Ecology and Conservation, 62, e03809. 10.1016/j.gecco.2025.e03809

Coop, G., Witonsky, D., Di Rienzo, A., & Pritchard, J. K. (2010). Using environmental correlations to identify loci underlying local adaptation. Genetics, 185(4), 1411–1423. 10.1534/genetics.110.114819

Czech, L., Spence, J. P., & Exposito-Alonso, M. (2024). grenedalf: population genetic statistics for the next generation of pool sequencing. Bioinformatics, 40(8). 10.1093/bioinformatics/btae508

Dam, H. G. (2013). Evolutionary adaptation of marine zooplankton to global change. Annual review of marine science, 5, 349–370. 10.1146/annurev-marine-121211-172229

Dam, H. G., deMayo, J. A., Park, G., Norton, L., He, X., Finiguerra, M. B., Baumann, H., Brennan, R. S., & Pespeni, M. H. (2021). Rapid, but limited, zooplankton adaptation to simultaneous warming and acidification. Nature Climate Change, 11(9), 780–786. 10.1038/s41558-021-01131-5

Dauphin, B., Rellstab, C., Wüest, R. O., Karger, D. N., Holderegger, R., Gugerli, F., & Manel, S. (2023). Re-thinking the environment in landscape genomics. Trends in ecology & evolution, 38(3), 261–274. 10.1016/j.tree.2022.10.010

Day, T., Dawe, C., Dobson, T., & Hillier, P. (1983). A chromosomal inversion polymorphism in Scandinavian populations of the seaweed fly, *Coelopa frigida*. Hereditas, 99(1), 135–145. https://onlinelibrary.wiley.com/doi/pdfdirect/10.1111/j.1601-5223.1983.tb00738.x?download=true

Delandmeter, P., & van Sebille, E. (2019). The Parcels v2.0 Lagrangian framework: new field interpolation schemes. Geosci. Model Dev., 12(8), 3571–3584. 10.5194/gmd-12-3571-2019

Diaz, J. A., Stern, D. B., & Lee, C. E. (2026). Local Adaptation Despite Gene Flow in Copepod Populations Across Salinity and Temperature Gradients in the Baltic Sea and North Sea Region. Diversity and Distributions, 32(5). 10.1111/ddi.70176

Dildar, T., Cui, W., Ikhwanuddin, M., & Ma, H. (2025). Aquatic Organisms in Response to Salinity Stress: Ecological Impacts, Adaptive Mechanisms, and Resilience Strategies. Biology (Basel), 14(6). 10.3390/biology14060667

Drushka, K., Asher, W. E., Ward, B., & Walesby, K. (2016). Understanding the formation and evolution of rain-formed fresh lenses at the ocean surface. Journal of Geophysical Research: Oceans, 121(4), 2673–2689. 10.1002/2015JC011527

Du, Y., Zhang, Y., & Shi, J. (2019). Relationship between sea surface salinity and ocean circulation and climate change. Science China Earth Sciences, 62(5), 771–782. 10.1007/s11430-018-9276-6

Du, Z., Wirtz, J., Zhou, Y. J., Jenstead, A., Opgenorth, T., Puls, A., Meyer, C., Gelembiuk, G. W., & Lee, C. E. (2025). Genome architecture evolution in an invasive copepod species complex. Nature communications, 16(1), 10312. 10.1038/s41467-025-65292-z

Durack, P. J., Wijffels, S. E., & Matear, R. J. (2012). Ocean salinities reveal strong global water cycle intensification during 1950 to 2000. Science, 336(6080), 455–458. 10.1126/science.1212222

Dutz, J., Zettler, M., Kremp, A., Paul, C., & Kube, S. (2025). Biological assessment of the Baltic Sea 2023 Meereswissenschaftliche Berichte Warnemünde, 129. 10.12754/msr-2025-0129

Ewels, P., Magnusson, M., Lundin, S., & Kaller, M. (2016). MultiQC: summarize analysis results for multiple tools and samples in a single report. Bioinformatics, 32(19), 3047–3048. 10.1093/bioinformatics/btw354

Farhadi, A., Liu, Y., Xu, C., Wang, X., & Li, E. (2022). The role of the renin-angiotensin system (RAS) in salinity adaptation in Pacific white shrimp (Litopenaeus vannamei) [Original Research]. Frontiers in Endocrinology, Volume 13 - 2022. 10.3389/fendo.2022.1089419

Fitzpatrick, M. C., & Keller, S. R. (2015). Ecological genomics meets community-level modelling of biodiversity: mapping the genomic landscape of current and future environmental adaptation. Ecol Lett, 18(1), 1–16. 10.1111/ele.12376

Fuhrmann, N., Prakash, C., & Kaiser, T. S. (2023). Polygenic adaptation from standing genetic variation allows rapid ecotype formation. Elife, 12. 10.7554/eLife.82824

Gain, C., & Francois, O. (2021). LEA 3: Factor models in population genetics and ecological genomics with R. Mol Ecol Resour, 21(8), 2738–2748. 10.1111/1755-0998.13366

Gain, C., Rhone, B., Cubry, P., Salazar, I., Forbes, F., Vigouroux, Y., Jay, F., & Francois, O. (2023). A Quantitative Theory for Genomic Offset Statistics. Mol Biol Evol, 40(6). 10.1093/molbev/msad140

Gassen, L., Esters, L., Ribas-Ribas, M., & Wurl, O. (2024). The impact of rainfall on the sea surface salinity: a mesocosm study. Scientific reports, 14(1), 6353. 10.1038/s41598-024-56915-4

Gautier, M. (2015). BayPass Genome-Wide Scan for Adaptive Differentiation and Association Analysis with population-specific covariables In

Gautier, M., Vitalis, R., Flori, L., & Estoup, A. (2022). f-Statistics estimation and admixture graph construction with Pool-Seq or allele count data using the R package poolfstat. Mol Ecol Resour, 22(4), 1394–1416. 10.1111/1755-0998.13557

Giallongo, G., Douek, J., Harbuzov, Z., Galil, B. S., & Rinkevich, B. (2021). Long-term changes in population genetic features of a rapidly expanding marine invader: implication for invasion success. Biological Invasions, 23(8), 2541–2552. 10.1007/s10530-021-02521-8

Hahn, A., Nascimento-Schulze, J. C., Avgerinou, G., Bayer, T., & Brennan, R. S. (2026). Local Adaptation and Osmoregulatory Mechanisms of the Copepod *Acartia tonsa* Under Low Salinity Stress. Mol Ecol, 35(1), e70208. 10.1111/mec.70208

Haldane, J. B. S. (1930). Theoretical genetics of autopolyploids. Journal of Genetics, 22(3), 359–372. 10.1007/bf02984197

Harianto, J., Carey, N., Byrne, M., & Price, S. (2019). respR—An R package for the manipulation and analysis of respirometry data. Methods in Ecology and Evolution, 10(6), 912–920. 10.1111/2041-210x.13162

Höper, A., Funk, N., Mittermayer, F., Temming, A., & Funk, S. (2025). Food intake of early juvenile western Baltic cod (*Gadus morhua*) during settlement transition. J Fish Biol. 10.1111/jfb.70234

Huang, Q., Xu, L., Xie, L., Liu, P., Rizo, E. Z. C., & Han, B. P. (2022). Spatial and temporal variation of genetic diversity and genetic differentiation in *Daphnia galeata* populations in four large reservoirs in southern China. Front Microbiol, 13, 1041011. 10.3389/fmicb.2022.1041011

Huang, S., Zhou, F., Yang, Q., Jiang, S., Chen, J., Xiong, J., Li, E., & Li, Y. (2026). Integrative Transcriptomic and Network Analysis of Hemocyte Volume Plasticity and Redox Regulation Under Osmotic Stress in Penaeus monodon. Antioxidants (Basel), 15(1). 10.3390/antiox15010147

Jain, K., & Devi, A. (2018). Polygenic adaptation in changing environments(a). EPL (Europhysics Letters), 123(4). 10.1209/0295-5075/123/48002

Jeffreys, H. (1961). Theory of Probability. In (3rd Edition ed., pp. 95–103). Oxford University Press.

Johannesson, K., Le Moan, A., Perini, S., & Andre, C. (2020). A Darwinian Laboratory of Multiple Contact Zones. Trends Ecol Evol, 35(11), 1021–1036. 10.1016/j.tree.2020.07.015

Jun, G., Wing, M. K., Abecasis, G. R., & Kang, H. M. (2015). An efficient and scalable analysis framework for variant extraction and refinement from population-scale DNA sequence data. Genome Res, 25(6), 918–925. 10.1101/gr.176552.114

Kankaanpää, H. T., Alenius, P., Kotilainen, P., & Roiha, P. (2023). Decreased surface and bottom salinity and elevated bottom temperature in the Northern Baltic Sea over the past six decades. Sci Total Environ, 859(Pt 2), 160241. 10.1016/j.scitotenv.2022.160241

Kawecki, T. J., & Ebert, D. (2004). Conceptual issues in local adaptation. Ecology letters, 7(12), 1225–1241. 10.1111/j.1461-0248.2004.00684.x

Kiørboe, T., Møhlenberg, F., & Hamburger, K. (1985). Bioenergetics of the planktonic copepod *Acartia tonsa*: relation between feeding, egg production and respiration, and composition of specific dynamic action. Marine Ecology Progress Series, 26, 85–97. 10.3354/meps026085

Kirkpatrick, M., & Barton, N. (2006). Chromosome inversions, local adaptation and speciation. Genetics, 173(1), 419–434. 10.1534/genetics.105.047985

Kniebusch, M., Meier, H. E. M., & Radtke, H. (2019). Changing Salinity Gradients in the Baltic Sea As a Consequence of Altered Freshwater Budgets. Geophysical Research Letters, 46(16), 9739–9747. 10.1029/2019gl083902

Kopp, M., & Hermisson, J. (2007). Adaptation of a quantitative trait to a moving optimum. Genetics, 176(1), 715–719. 10.1534/genetics.106.067215

Lance, J. (1962). Effects of water of reduced salinity on the vertical migration of zooplankton. Journal of the Marine Biological Association of the United Kingdom, 42(2), 131–154. 10.1017/s0025315400001272

Lance, J. (1965). Respiration and Osmotic Behaviour of the Copepod *Acartia tonsa* in Diluted Sea Water. Comp Biochem Physiol, 14, 155–165. 10.1016/0010-406x(65)90016-2

Le Moan, A., Bekkevold, D., & Hemmer-Hansen, J. (2021). Evolution at two time frames: ancient structural variants involved in post-glacial divergence of the European plaice (*Pleuronectes platessa*). Heredity (Edinb), 126(4), 668–683. 10.1038/s41437-020-00389-3

Leandro, S. M., Tiselius, P., & Queiroga, H. (2006). Growth and development of nauplii and copepodites of the estuarine copepod Acartia tonsa from southern Europe (Ria de Aveiro, Portugal) under saturating food conditions. Marine Biology, 150(1), 121–129. 10.1007/s00227-006-0336-y

Lee, C. E. (2021). Ion Transporter Gene Families as Physiological Targets of Natural Selection During Salinity Transitions in a Copepod. Physiology (Bethesda), 36(6), 335–349. 10.1152/physiol.00009.2021

Lee, C. E. (2023). Genome architecture underlying salinity adaptation in the invasive copepod *Eurytemora affinis* species complex: A review. iScience, 26(10), 107851. 10.1016/j.isci.2023.107851

Lee, C. E., Downey, K., Colby, R. S., Freire, C. A., Nichols, S., Burgess, M. N., & Judy, K. J. (2022). Recognizing Salinity Threats in the Climate Crisis. Integr Comp Biol, 62(2), 441–460. 10.1093/icb/icac069

Lenth, R. V. (2023). emmeans: Estiamte Marginal Means, aka Least-Squares Means. In (Version R package version 1.8.4-1) https://CRAN.R-project.org/package=emmeans

Li, H., Handsaker, B., Wysoker, A., Fennell, T., Ruan, J., Homer, N., Marth, G., Abecasis, G., Durbin, R., & Genome Project Data Processing, S. (2009). The Sequence Alignment/Map format and SAMtools. Bioinformatics, 25(16), 2078–2079. 10.1093/bioinformatics/btp352

Li, Y.-D., Si, M.-R., Jiang, S.-G., Yang, Q.-B., Jiang, S., Yang, L.-S., Huang, J.-H., Chen, X., Zhou, F.-L., & Li, E. (2023). Transcriptome and molecular regulatory mechanisms analysis of gills in the black tiger shrimp Penaeus monodon under chronic low-salinity stress [Original Research]. Frontiers in Physiology, Volume 14–2023. 10.3389/fphys.2023.1118341

Lovell, R. S. L., Collins, S., Martin, S. H., Pigot, A. L., & Phillimore, A. B. (2023). Space-for-time substitutions in climate change ecology and evolution. Biol Rev Camb Philos Soc, 98(6), 2243–2270. 10.1111/brv.13004

Luu, K., Bazin, E., & Blum, M. G. (2017). pcadapt: an R package to perform genome scans for selection based on principal component analysis. Mol Ecol Resour, 17(1), 67–77. 10.1111/1755-0998.12592

Magouz, F. I., Essa, M. A., Matter, M., Mansour, A. T., Gaber, A., & Ashour, M. (2021). Effect of Different Salinity Levels on Population Dynamics and Growth of the Cyclopoid Copepod *Oithona nana*. Diversity, 13(5). 10.3390/d13050190

Mantel, L. H., & Farmer, L. L. (1983). Osmotic and ionic regulation. In Internal anatomy and physiological regulation (Vol. 5, pp. 53–161).

Matschiner, M., Barth, J. M. I., Torresen, O. K., Star, B., Baalsrud, H. T., Brieuc, M. S. O., Pampoulie, C., Bradbury, I., Jakobsen, K. S., & Jentoft, S. (2022). Supergene origin and maintenance in Atlantic cod. Nat Ecol Evol, 6(4), 469–481. 10.1038/s41559-022-01661-x

Mauchline, J. (1998). The Biology of Calanoid Copepods (Vol. 33). Academic Press.

Nour, O. M., Pansch, C., Lenz, M., Wahl, M., Clemmesen, C., & Stumpp, M. (2021). Impaired larval development at low salinities could limit the spread of the non-native crab *Hemigrapsus takanoi* in the Baltic Sea. Aquatic Biology, 30, 85–99.

Okonechnikov, K., Conesa, A., & García-Alcalde, F. (2015). Qualimap 2: advanced multi-sample quality control for high-throughput sequencing data. Bioinformatics, 32(2), 292–294. 10.1093/bioinformatics/btv566

Pante, E., Simon-Bouhet, B., & Irisson, J. (2023). marmap: Import, Plot and Analyze Bathymetric and Topographic Data. In (Version R package version 1.0.10) https://CRAN.R-project.org/package=marmap

Pfrender, M. E., & Lynch, M. (2000). Quantitative genetic variation in *Daphnia*: temporal changes in genetic architecture. Evolution, 54(5), 1502–1509. 10.1111/j.0014-3820.2000.tb00696.x

Pinseel, E., Ruck, E. C., Nakov, T., Jonsson, P. R., Kourtchenko, O., Kremp, A., Pinder, M. I. M., Roberts, W. R., Sjoqvist, C., Topel, M., Godhe, A., Hahn, M. W., & Alverson, A. J. (2025). Genome-Wide Adaptation to a Complex Environmental Gradient in a Keystone Phytoplankton Species. Mol Ecol, 34(13), e17817. 10.1111/mec.17817

Posavi, M., Gelembiuk, G. W., Larget, B., & Lee, C. E. (2014). Testing for beneficial reversal of dominance during salinity shifts in the invasive copepod Eurytemora affinis, and implications for the maintenance of genetic variation. Evolution, 68(11), 3166–3183. 10.1111/evo.12502

Posavi, M., Gulisija, D., Munro, J. B., Silva, J. C., & Lee, C. E. (2020). Rapid evolution of genome-wide gene expression and plasticity during saline to freshwater invasions by the copepod *Eurytemora affinis* species complex. Mol Ecol, 29(24), 4835–4856. 10.1111/mec.15681

Quinlan, A. R., & Hall, I. M. (2010). BEDTools: a flexible suite of utilities for comparing genomic features. Bioinformatics, 26(6), 841–842. 10.1093/bioinformatics/btq033

Redeke, H. C. (1934). On the Occurrence of two Pelagic Copepods, *Acartia bifilosa* and *Acartia tonsa*, in the Brackish Waters of the Netherlands. ICES Journal of Marine Science, 9(1), 39–45. 10.1093/icesjms/9.1.39

Rellstab, C., Dauphin, B., & Exposito-Alonso, M. (2021). Prospects and limitations of genomic offset in conservation management. Evol Appl, 14(5), 1202–1212. 10.1111/eva.13205

Reusch, T. B. H., Dierking, J., Andersson, H. C., Bonsdorff, E., Carstensen, J., Casini, M., Czajkowski, M., Hasler, B., Hinsby, K., Hyytiäinen, K., Johannesson, K., Jomaa, S., Jormalainen, V., Kuosa, H., Kurland, S., Laikre, L., MacKenzie, B. R., Margonski, P., Melzner, F.,…Zandersen, M. (2018). The Baltic Sea as a time machine for the future coastal ocean. Science advances, 4(5), eaar8195. 10.1126/sciadv.aar8195

Rueda Moreno, G., & Sasaki, M. C. (2023). Starvation reduces thermal limits of the widespread copepod Acartia tonsa. Ecol Evol, 13(10), e10586. 10.1002/ece3.10586

Sanford, E., & Kelly, M. W. (2011). Local adaptation in marine invertebrates. Annual review of marine science, 3, 509–535. 10.1146/annurev-marine-120709-142756

Schneider, C. A., Rasband, W. S., & Eliceiri, K. W. (2012). NIH Image to ImageJ: 25 years of image analysis. Nature methods, 9(7), 671–675. 10.1038/nmeth.2089

Serandour, B., Blenckner, T., Jan, K. M. G., Leroy, B., Ramiro-Sánchez, B., Campbell, E., & Winder, M. (2024). Spatial distribution projections of suitable environmental conditions for key Baltic Sea zooplankton species. Limnology and Oceanography, 69(12), 2801–2814. 10.1002/lno.12705

Seuront, L. (2006). Effect of salinity on the swimming behaviour of the estuarine calanoid copepod Eurytemora affinis. Journal of Plankton Research, 28(9), 805–813.

Snoeijs-Leijonmalm, P., Schubert, H., & Radziejewska, T. (2017). Biological Oceanography of the Baltic Sea (P. Snoeijs-Leijonmalm, H. Schubert, & T. Radziejewska, Eds. 1 ed.). Spriinger Dordrecht. 10.1007/978-94-007-0668-2

Stern, D. B., Anderson, N. W., Diaz, J. A., & Lee, C. E. (2022). Genome-wide signatures of synergistic epistasis during parallel adaptation in a Baltic Sea copepod. Nat Commun, 13(1), 4024. 10.1038/s41467-022-31622-8

Stern, D. B., & Lee, C. E. (2020). Evolutionary origins of genomic adaptations in an invasive copepod. Nat Ecol Evol, 4(8), 1084–1094. 10.1038/s41559-020-1201-y

Storey, J. D., Bass, A. J., Dabney, A., & Robinson, D. (2022). qvalue: Q-value estimation for false discovery rate control. In (Version v.2.30.0) http://github.com/jdstorey/qvalue

Svetlichny, L., & Hubareva, E. (2014). Salinity tolerance of alien copepods *Acartia tonsa* and *Oithona davisae* in the Black Sea. Journal of Experimental Marine Biology and Ecology, 461, 201–208. 10.1016/j.jembe.2014.08.012

Team, R. C. (2022). R: A language and environment for statistical computing. In (Version 4.2.2)

Tigano, A., & Friesen, V. L. (2016). Genomics of local adaptation with gene flow. Mol Ecol, 25(10), 2144–2164. 10.1111/mec.13606

Turner, J. T. (1984). The feeding ecology of some zooplankters that are important prey items of larval fish.

Vasimuddin, M., Misra, S., H., L., & S., A. (2019, 20-24 May 2019). Efficient Architecture-Aware Acceleration of BWA-MEM for Multicore Systems. 2019 IEEE International Parallel and Distributed Processing Symposium (IPDPS),

Walters, R. J., & Berger, D. (2019). Implications of existing local (mal)adaptations for ecological forecasting under environmental change. Evol Appl, 12(7), 1487–1502. 10.1111/eva.12840

Wellenreuther, M., Merot, C., Berdan, E., & Bernatchez, L. (2019). Going beyond SNPs: The role of structural genomic variants in adaptive evolution and species diversification. Mol Ecol, 28(6), 1203–1209. 10.1111/mec.15066

Wellenreuther, M., Oomen, R. A., Han, K. Y., Krohman, R., & Reusch, T. B. H. (2025). Beyond supergenes: the diverse roles of inversions in trait evolution. Trends in ecology & evolution, 40(11), 1054–1065. 10.1016/j.tree.2025.08.004

Witalis, B., Iglikowska, A., Ronowicz, M., Weydmann-Zwolicka, A., & Kukliński, P. (2024). The zooplankton community of Baltic Sea ports: diversity and seasonal dynamics. Oceanologia. 10.1016/j.oceano.2024.02.001

Wu, T., Hu, E., Xu, S., Chen, M., Guo, P., Dai, Z., Feng, T., Zhou, L., Tang, W., Zhan, L., Fu, X., Liu, S., Bo, X., & Yu, G. (2021). clusterProfiler 4.0: A universal enrichment tool for interpreting omics data. Innovation (Camb), 2(3), 100141. 10.1016/j.xinn.2021.100141

Yeaman, S. (2015). Local Adaptation by Alleles of Small Effect. Am Nat, 186 *Suppl 1*, S74–89. 10.1086/682405

Yeaman, S., & Whitlock, M. C. (2011). The genetic architecture of adaptation under migration-selection balance. Evolution, 65(7), 1897–1911. 10.1111/j.1558-5646.2011.01269.x

