## Supplements for "Salinity adaptation in a copepod reveals contrasting genomic architectures and the limits of genomic offsets"

**for**

### Affiliations


This file includes:

Supplementary Methods – connectivity

Supplementary Discussion – candidate genes

Supplementary Tables S1 – S3 (link provided to large tables S4 – S6)

Supplementary Figures S1 – S12

### SUPPLEMENTARY METHODS

#### Connectivity analysis

*Acartia tonsa* begin to emerge in early summer from overwintering eggs in the sediment, triggered by increasing temperatures (Sullivan et al., 2007). They reproduce in the upper water column until temperatures drop below 10-12°C (Holste & Peck, 2006), when they resume producing resting eggs and enter diapause. In the Baltic Sea, this results in *A. tonsa* occurring in the plankton from approximately May to November (Katajisto et al., 1998; Witalis et al., 2024; Dutz et al., 2025). Our simulations only cover the core reproductive months from June to October to ensure we are capturing realistic spawning conditions at all locations. The maximum dispersal duration was set to 28 days which represents the time from hatching to the end of the reproductive period of *A. tonsa,* considering that larvae reach maturity approximately 10 days after hatching (Parrish & Wilson, 1978) and the reproductive time lasts around 20 days *(Shayegan et al., 2016)*. Since *in-situ* swimming behavior and velocities are not fully characterized for *A. tonsa*, we assumed copepods disperse within the upper 25 m of the water column, corresponding to the depth range covered during diel migrations (Holliland et al., 2012). Note that due to the discrete numerical grid on which we prescribed the particle velocities, for some near-shore release sites not all of the 1000 drifters released per day and release site, actually drifted freely. We accounted for this, by adapting N per site to the actual number of freely drifting particles.

### SUPPLEMENTARY DISCUSSION

#### Candidate regions of North Sea-Baltic adaptation are linked to salinity tolerance

When looking into the functions of genes associated with putatively adaptive loci, two genomic regions related to the North Sea– Baltic Sea contrast are worth highlighting. First, the region on chromosome 8 that was characterized by the strongest peak in $F_{ST}$ contained genes linked to acute stress responses and homeostasis with several key osmoregulatory genes: The cytochrome P450 superfamily has been found to respond to stress in copepods and other marine invertebrates (Snyder, 2000; Lauritano et al., 2012) and genes belonging to the same family are linked to salinity tolerance in plants (Krishnamurthy et al., 2020). In the Chinese mitten crab *Eriocheir sinensis* cytochrome P450 has been upregulated in saltwater acclimated crabs compared to freshwater individuals (Yang et al., 2022). Our previous work comparing North Sea and Baltic Sea populations found that glycine transport was downregulated in response to hypo-osmotic stress (Hahn et al., 2026). Glycine is an important osmolyte in many marine invertebrates (Podbielski et al., 2022) and glycine receptors have been linked to maintaining homeostasis under stress in marine pteropods (Moya et al., 2016). Here, we found the glycine receptor subunit a-4 was under selection on chromosome 8, further supporting its role in salinity homeostasis and adaptation. Lastly, the neuronal calcium sensor-2 was under selection and has been linked to ion transport and was upregulated in the gills of Chinese mitten crabs exposed to high salinity (Yang et al., 2022). Together the genes on the chromosome 8 region of interest appear to be important for responding to salinity stress, potentially allowing *A. tonsa* to tolerate hypo-osmotic conditions by supporting cellular homeostasis.

Second, the chromosome 6 region contained one gene directly related to transmembrane transport as well as several genes involved in transcriptional and epigenetic regulation. SLC71a1 belongs to a superfamily of solute carrying proteins that are known to facilitate transmembrane transport of a broad range of molecules from sugars, to ions and amino acids (Pizzagalli et al., 2021). While there is no extensive record of the function of this specific solute carrier, members of the same superfamily (SLC12) linked to salinity stress in a gene expression study on ridgetail white shrimp (Tang et al., 2025), and have been differentially expressed in *A. tonsa* populations from North Sea and Baltic Sea exposed to low salinity stress (Hahn et al., 2026). Thus, the presence of salinity-related genes within the islands of divergence may highlight their role in facilitating the adaptation to the low-salinity Baltic.

### GEA loci are linked to osmoregulation

We observed a large overlap between loci corresponding to salinity range and mean salinity at our sampling sites. This indicates that similar gene responses facilitate adaptation to absolute salinity but also the variability of the environment. In our system, mean salinity was correlated with the overall variability, with Western Baltic regions strongly influenced by North Sea inflow events and therefore characterized by both higher and more variable salinity levels compared to stable Eastern Baltic regions. Therefore, there was a large overlap in loci and genes identified correlated with both variables. In addition, a considerable number of genes were correlated with the environment and also identified as targets of strong directional selection. Among these were key genes that have been linked to ion regulatory processes of animals experiencing hypoosmotic stress such as ion transporters or exchangers. In crustaceans, an interplay of $Na^{+}/H^{+}$-exchangers or antiporters (NHEs and NHAs), and $Na^{+}/K^{+}$-ATPases (NKAs) has been proposed to facilitate sodium uptake against a strong osmotic gradients, e.g., in freshwater habitats (Lee et al., 2022). We identified these key genes alongside other osmoregulatory relevant genes, highlighting that *A. tonsa* uses similar pathways when adapting to low salinity and fluctuating habitats.

### SUPPLEMENTARY TABLES

**Table S1 Overview of sampling locations**. Sea surface mean salinity and salinity range in practical salinity units (PSU) calculated from CMEMS (https://doi.org/10.48670/moi-00021). For each sampling station (Fig. 1A) salinity was calculated from June to October for the sampling year plus the five years prior to capture per-site variability, values include standard deviation. For SW08 where the same station was sampled twice (within less than 1 $km^2$), we used the exact coordinates for the first sampling but calculated salinity values for both sampling times. The colors used for each location are consistent throughout the manuscript.

| Location | Longitude | Latitude | Sampling date | Mean salinity | Salinity range | Culture | Genomics |
|---|---|---|---|---|---|---|---|
| WH | 8.1490 | 53.5130 | 2022-09-09 | 25.3 ± 1.1 | 6.2 | Yes | Yes |
| SK | 8.9487 | 54.0929 | 2022-08-21 | 27.3 ± 1.5 | 8.8 | Yes | Yes |
| BOK | 10.034 | 54.5169 | 2023-09-09 | 13.2 ± 2.0 | 10.0 | Yes | Yes |
| KB03 | 10.1533 | 54.6884 | 2022-08-30 | 13.1 ± 2.0 | 10.2 | Yes | Yes |
| SW08 | 11.6161 | 54.4079 | 2022-08-31<br>2023-08-31 | 10.9 ± 2.0<br>11.1 ± 1.9 | 11.2<br>10.6 | Yes (2022) | Yes (2022+2023) |
| H21 | 13.4914 | 54.9433 | 2023-09-01 | 8.1 ± 0.4 | 2.9 | Yes | Yes |
| BB23 | 15.7503 | 55.2912 | 2023-09-02 | 7.6 ± 0.2 | 1.1 | Yes | No |
| BB36 | 15.9981 | 54.9589 | 2022-09-07 | 7.6 ± 0.2 | 1.0 | No | Yes |
| GDY | 18.5642 | 54.5329 | 2023-09-15 | 6.9 ± 0.3 | 1.7 | Yes | Yes |
| GB96 | 19.5827 | 56.9165 | 2022-09-03 | 7.1 ± 0.1 | 0.7 | No | Yes |
| LL17 | 21.0478 | 59.0200 | 2023-08-15 | 6.4 ± 0.3 | 1.4 | Yes | No |
| IU7 | 21.2021 | 59.4890 | 2023-08-09 | 6.1 ± 0.2 | 0.9 | Yes | No |
| FIN | 23.2663 | 59.7766 | 2018-09-14 | 5.3 ± 0.3 | 1.3 | No | Yes |

**Table S2** Sample statistics for all 11 sequencing pools used in the analysis. QC results from merged, duplicate-marked, and realigned samples done in *flagstat* and *qualimap*.

| Station | Mapped reads | Mapped % | Properly paired % | Coverage mean | Coverage std | GC % | Mapping quality mean | Insert size median |
|---|---|---|---|---|---|---|---|---|
| BB36_22 | 490,071,468 | 88.12 | 68.19 | 68.0676 | 102.9036 | 32.61 | 36.2929 | 371.0 |
| BOK_23 | 661,546,708 | 92.02 | 71.72 | 93.0192 | 146.9587 | 32.39 | 36.0257 | 355.0 |
| FIN_18 | 519,804,966 | 83.62 | 64.56 | 72.6803 | 125.7872 | 32.6 | 36.2658 | 353.0 |

| | | | | | | | | |
|---|---|---|---|---|---|---|---|---|
| GB96_22 | 668,105,172 | 88.74 | 68.65 | 94.1608 | 136.4769 | 32.59 | 36.3994 | 361.0 |
| GDY_23 | 640,530,151 | 94.78 | 73.66 | 91.4543 | 119.5746 | 32.73 | 36.2061 | 356.0 |
| H21_23 | 538,777,757 | 93.55 | 72.23 | 76.6863 | 98.1196 | 32.47 | 36.4223 | 373.0 |
| KB03_22 | 477,241,153 | 94.41 | 72.70 | 68.821 | 92.452 | 32.62 | 36.3559 | 380.0 |
| SK_22 | 500,283,687 | 95.70 | 73.81 | 71.2342 | 135.5847 | 32.66 | 35.7116 | 371.0 |
| SW08_22 | 432,273,656 | 92.90 | 74.58 | 57.7856 | 89.2505 | 32.73 | 36.1938 | 342.0 |
| SW08_23 | 531,273,163 | 94.59 | 73.29 | 75.8309 | 93.9168 | 32.44 | 36.5638 | 362.0 |
| WH_22 | 622,868,255 | 95.30 | 73.84 | 88.7243 | 124.493 | 32.64 | 36.446 | 366.0 |

**Table S3** Number of SNPs and genes that are shared between two or more different methods. $GEA_{mean}$ = genotype-environment association for mean surface salinity, $GEA_{range}$ = genotype-environment association for salinity variability (lowest to highest value), $B_{select}$ = selection scan within the Baltic Sea, $NB_{select}$ = selection scan between North Sea and Baltic Sea.

| **Overlapping sets** | **SNP overlap** | **Gene overlap** |
|---|---|---|
| $GEA_{mean}$,$GEA_{range}$ | 1605 | 1577 |
| $GEA_{mean}$,$B_{select}$ | 24 | 264 |
| $GEA_{range}$,$B_{select}$ | 35 | 342 |
| $GEA_{mean}$,$NB_{select}$ | 0 | 165 |
| $GEA_{range}$,$NB_{select}$ | 1 | 204 |
| $GEA_{mean}$,$B_{select}$,$NB_{select}$ | 0 | 34 |
| $GEA_{range}$,$B_{select}$,$NB_{select}$ | 0 | 46 |

Additional large supplementary tables are archived on Zenodo:

https://doi.org/10.5281/zenodo.21358410

Table S4 Annotation of significant outlier genes.

Table S5 Enrichment of significant genes per analyses including KEGG and GO enrichment.

Table S6 Metadata and sample accessions of all WGR samples used in this study.

### SUPPLEMENTARY FIGURES

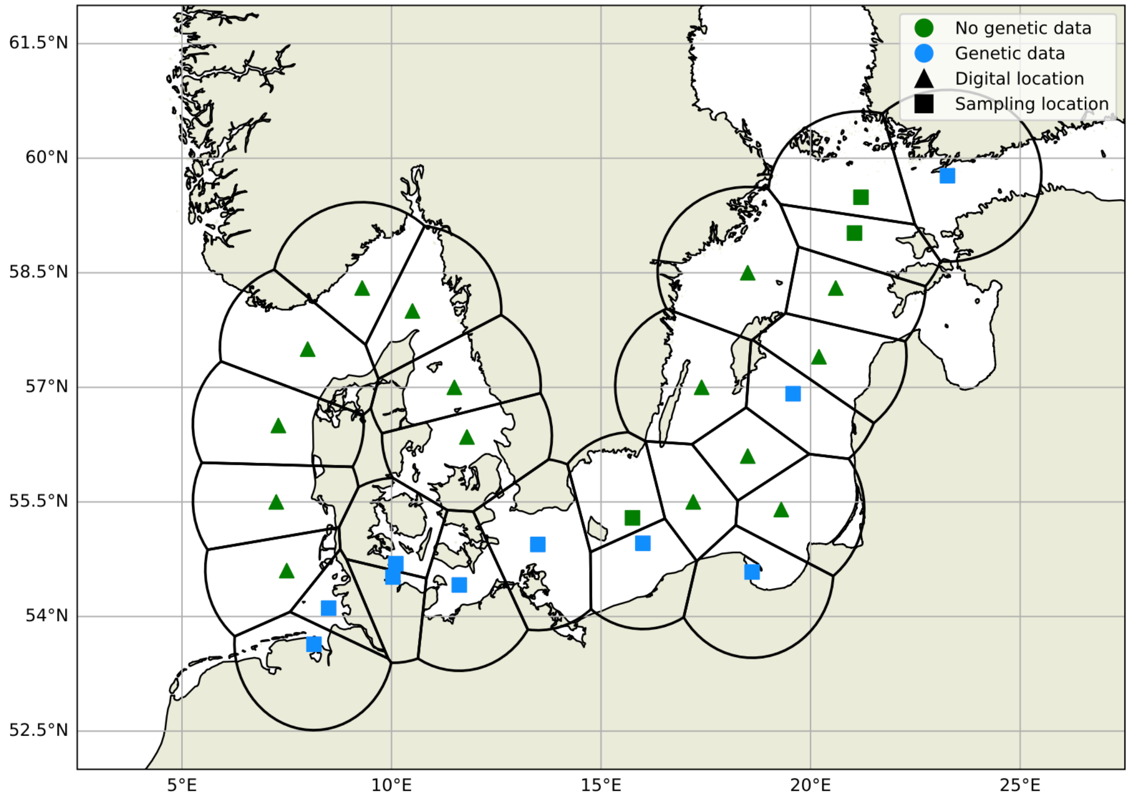


**Figure S1** Locations of sampling sites (squares) and additional intermediate sites (triangles) including Voronoi polygons used for the dispersal modeling. Genetic information is available for stations highlighted in blue.

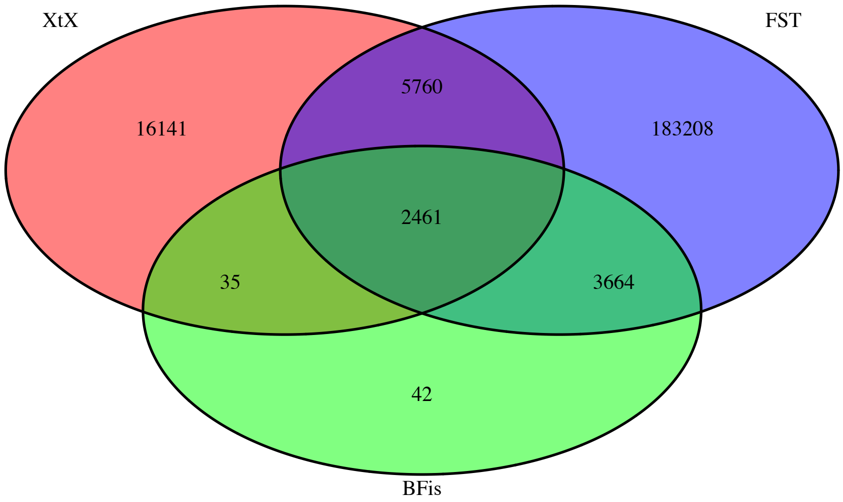


**Figure S2** Venn diagram of North Sea – Baltic Sea outlier analyses, green represent BFis > 20 dB, red represents calibrated XtX outliers, blue represents top 1% of pairwise $F_{ST}$.

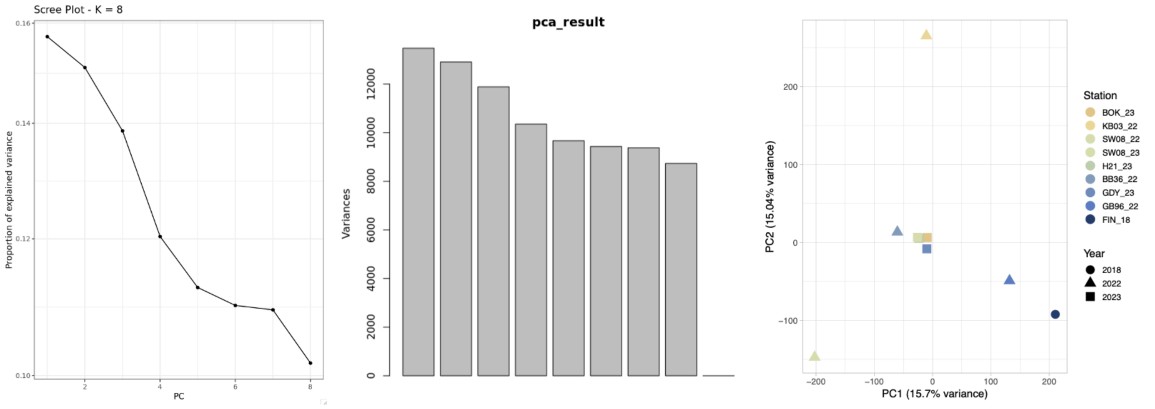


**Figure S3** Screeplot of and explained variance of Baltic allele frequency PCA. Based on the inflection point K = 4 was chosen for PCAdapt and LFMM.

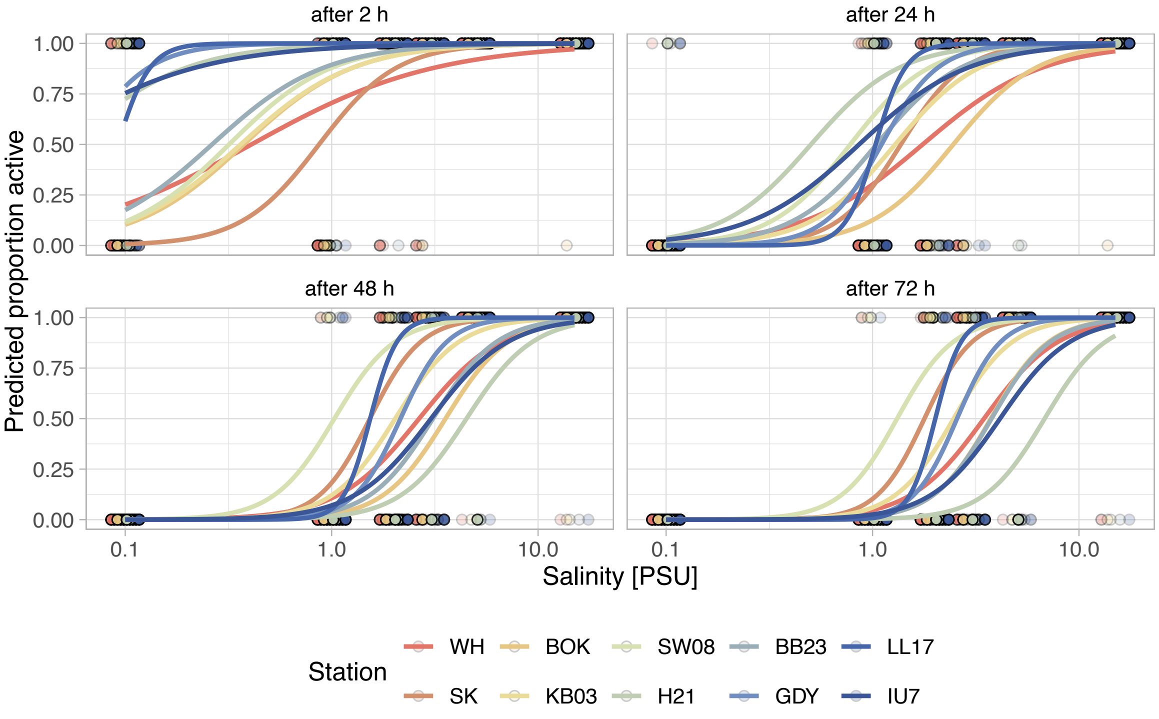


**Figure S4** Survival at low salinity measured 2, 24, 48, and 72 hours after exposure to treatment salinities, response curves are predicted from the *glmer*, colored points denote presence of active animals (0 = dead, 1 = alive), intensity denotes number of observed individuals.

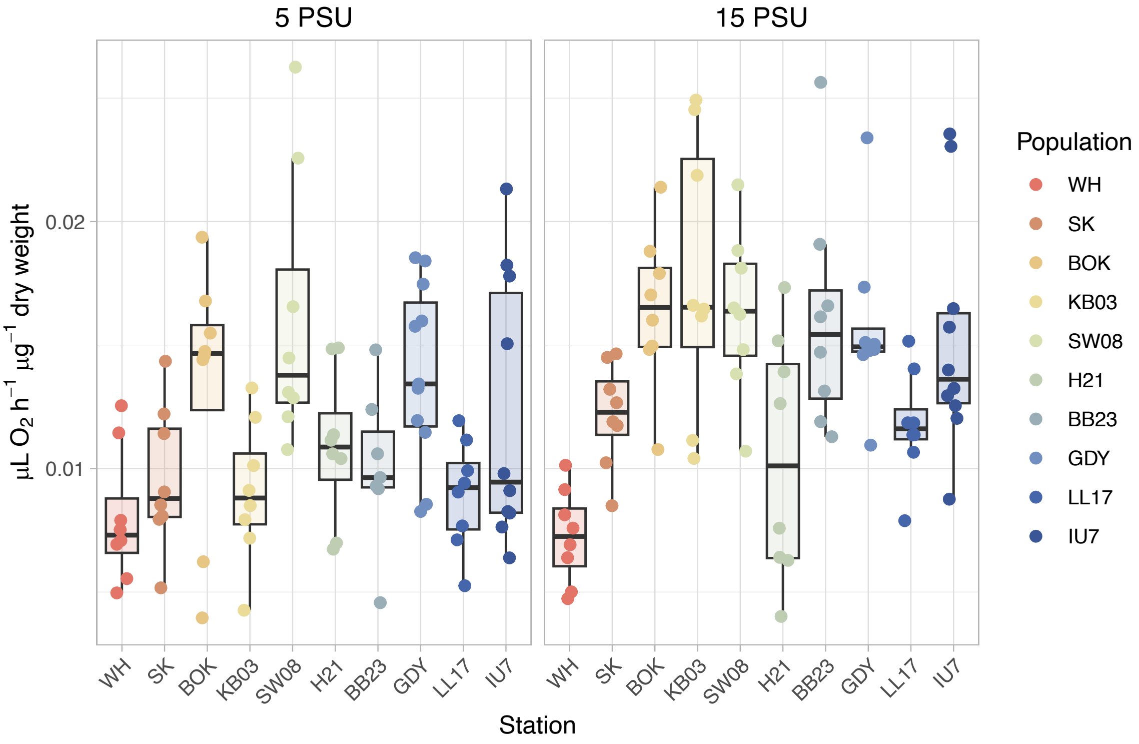


**Figure S5** Respiration rates of starved females (16 hours). Individual measurements are denoted by points. All rates are standardized by dry weight estimated from body size measurements.

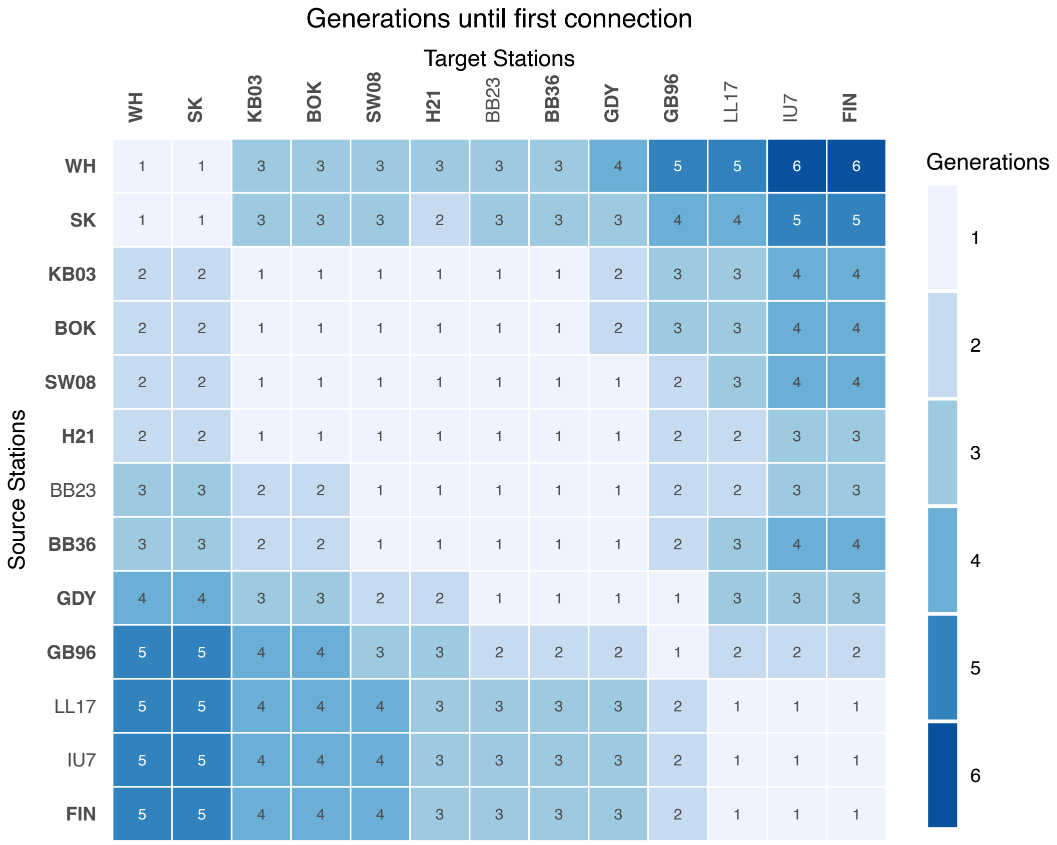


**Figure S6** Number of generations required to connect each station pair. Sampling sites with genetic data are highlighted in bold.

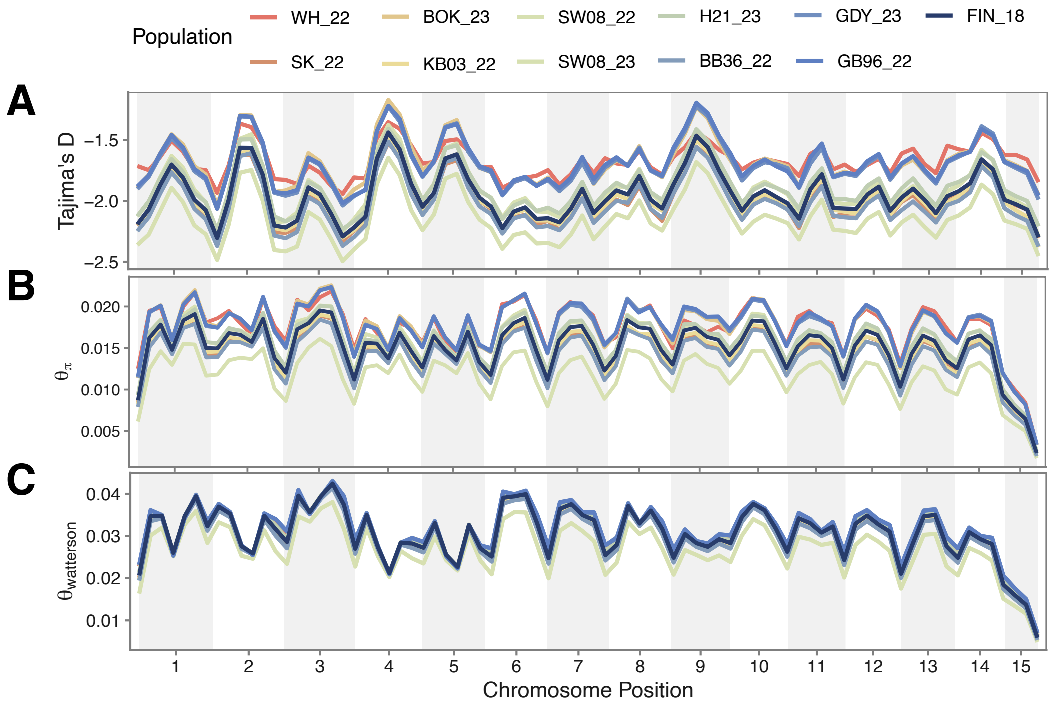


**Figure S7** Genome-wide population statistics estimated in 50 kb non-overlapping windows shown along the genome. A) Tajima's D B) Nucelotide diversity $\theta_\pi$ C) Watterson's $\theta$. All statistics were calculated in grenedalf. A smoothing function was applied in R for visual ease.

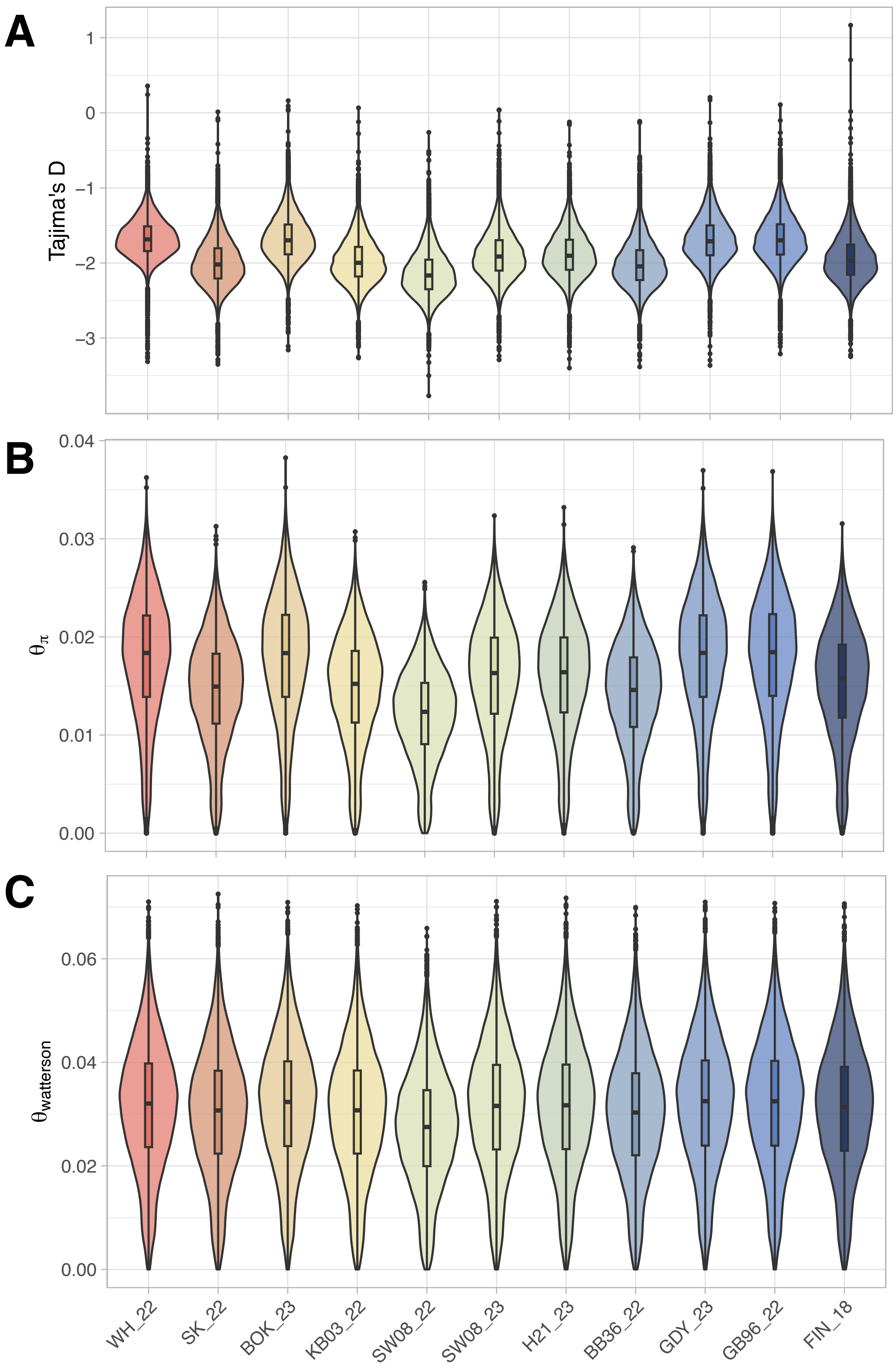


**Figure S8** Genome-wide population statistics estimated in 50 kb non-overlapping. A) Tajima's D B) Nucelotide diversity $\theta_\pi$ C) Watterson's $\theta$. All statistics were calculated in grenedalf.

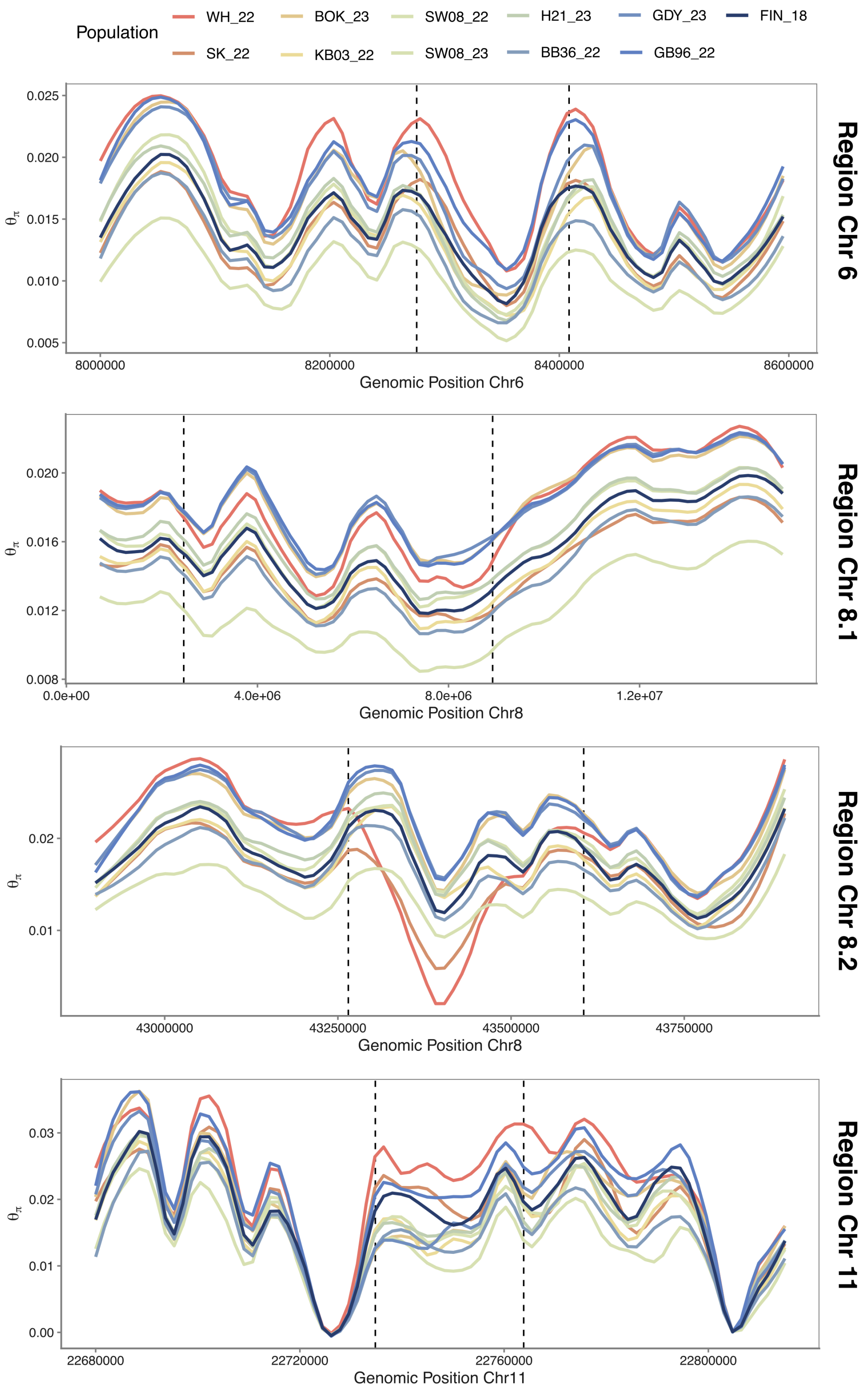


**Figure S9** Nucleotide diversity $\theta_\pi$ in 5 kb non-overlapping windows for regions of interest. Boundaries of SNP heavy outlier regions marked by dashed lines. A smoothing function was applied in R for visual ease.

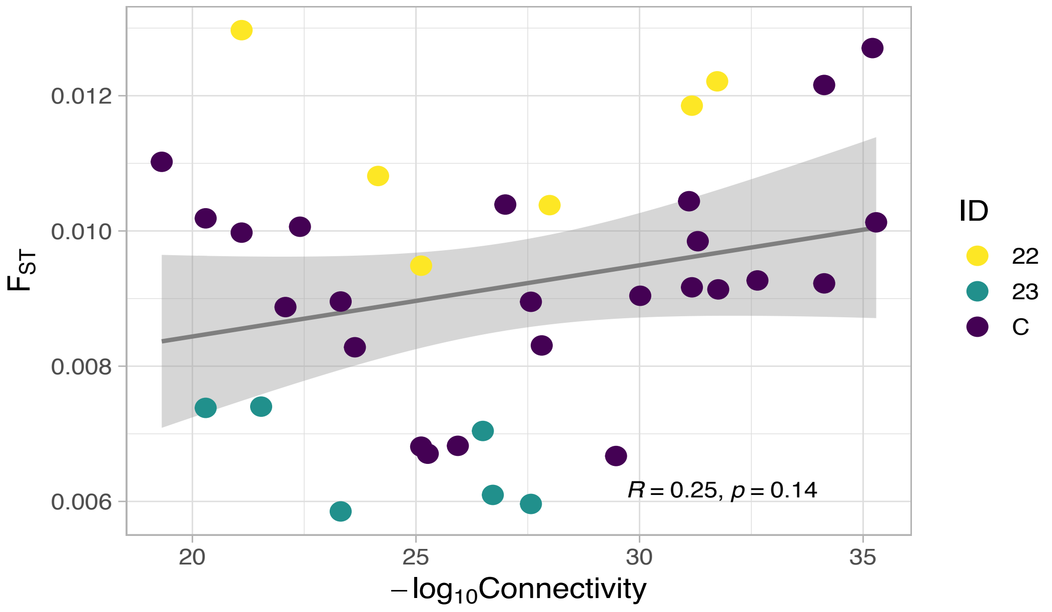


**Figure S10** Relationship of maximum connectivity and fixation index ($F_{ST}$). Colors are indicating sampling year, showing a strong batch effect regardless of connectivity.

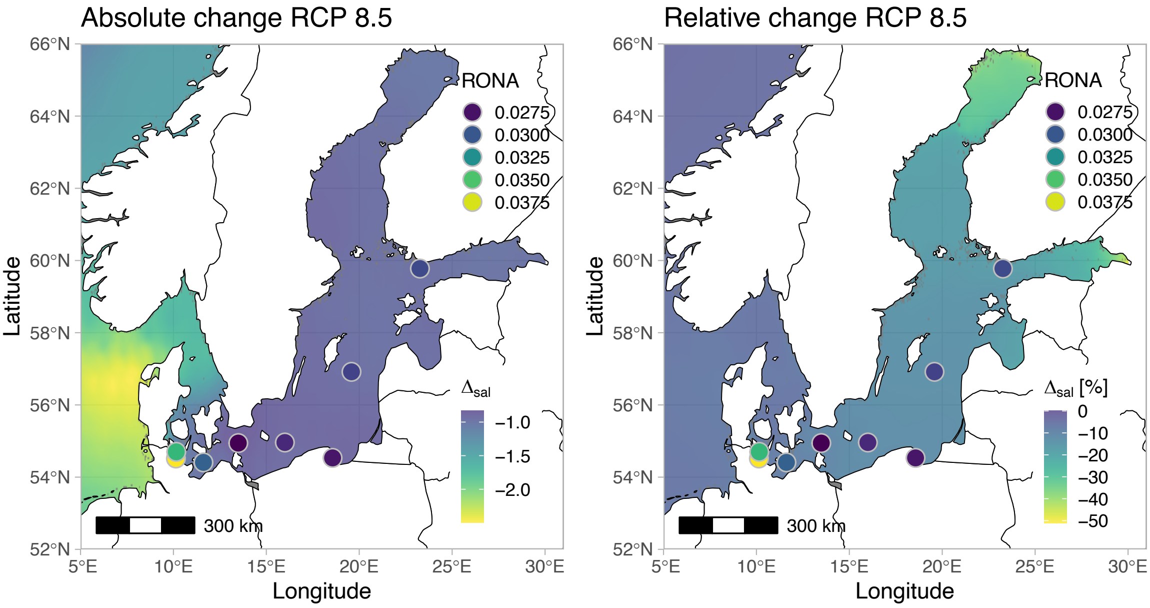


**Figure S11** Predicted salinity changes based on BioOracle 3 under the RCP 8.5 climate scenario overlaid with RONA (risk of non-adaptedness) predictions for the Baltic sampling locations. Higher RONA values indicate higher higher risk. Left: absolute predicted salinity change (in PSU). Right: relative predicted salinity change (in %).

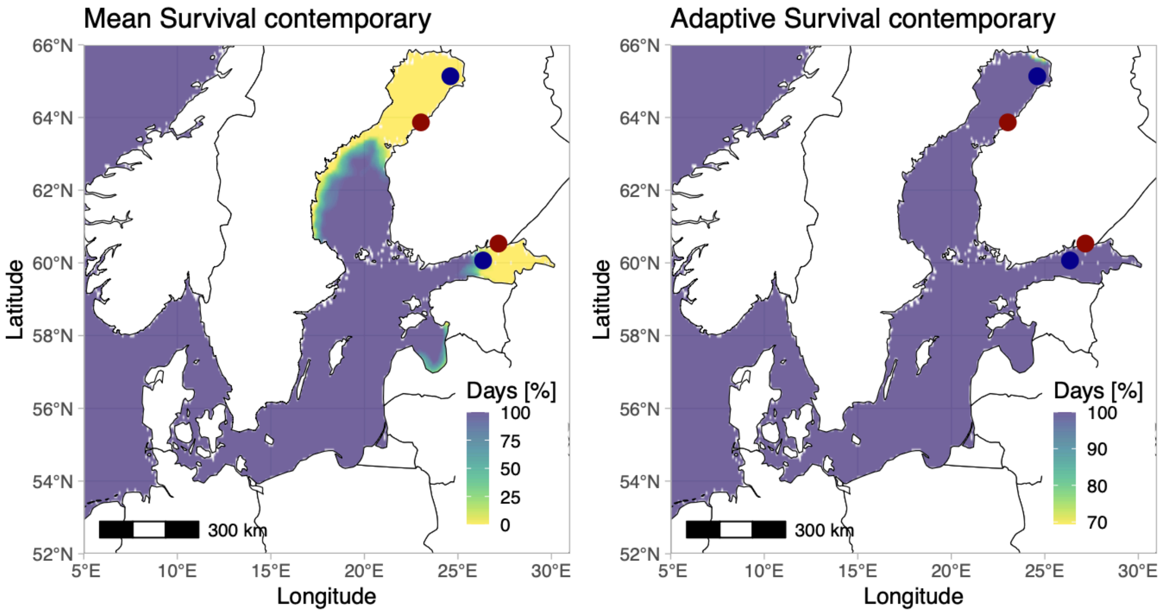


**Figure S12** Predicted contemporary habitat suitability based on experimentally determined thresholds (5% mortality after 24 hours), for species mean (left), and best performing population (right, LL17). Habitat suitability is shown as a percentage of days above the determined thresholds (0% = all days below viability threshold, 100% = all days above viability threshold), surface salinity data from CMEMS Global Ocean Physics Reanalysis is a daily mean averaged for the years 2020-2024. Dark blue circles indicate northern- and easternmost observation of *A. tonsa* in the OBIS database, dark red circles indicate the norther- and easternmost observation of *A. tonsa* in the HELCOM database.

### SUPPLEMENTARY REFERENCES


Dutz, J., Zettler, M., Kremp, A., Paul, C., & Kube, S. (2025). Biological assessment of the Baltic Sea 2023 *Meereswissenschaftliche Berichte Warnemünde*, *129*. https://doi.org/https://doi.org/10.12754/msr-2025-0129

Hahn, A., Nascimento-Schulze, J. C., Avgerinou, G., Bayer, T., & Brennan, R. S. (2026). Local Adaptation and Osmoregulatory Mechanisms of the Copepod *Acartia tonsa* Under Low Salinity Stress. *Mol Ecol*, *35*(1), e70208. https://doi.org/10.1111/mec.70208

Holliland, P. B., Ahlbeck, I., Westlund, E., & Hansson, S. (2012). Ontogenetic and seasonal changes in diel vertical migration amplitude of the calanoid copepods *Eurytemora affinis* and *Acartia spp.* in a coastal area of the northern Baltic proper. *Journal of Plankton Research*, *34*(4), 298-307. https://doi.org/10.1093/plankt/fbs001

Holste, L., & Peck, M. A. (2006). The effects of temperature and salinity on egg production and hatching success of Baltic *Acartia tonsa* (*Copepoda: Calanoida*): a laboratory investigation. *Marine Biology*, *148*(5), 1061–1070. https://doi.org/10.1007/s00227-005-0132-0

Katajisto, T., Viitasalo, M., & Koski, M. (1998). Seasonal occurrence and hatching of calanoid eggs in sediments of the northern Baltic Sea. *Marine Ecology Progress Series*, *163*, 133-143. https://doi.org/10.3354/meps163133

Krishnamurthy, P., Vishal, B., Ho, W. J., Lok, F. C. J., Lee, F. S. M., & Kumar, P. P. (2020). Regulation of a Cytochrome P450 Gene CYP94B1 by WRKY33 Transcription Factor Controls Apoplastic Barrier Formation in Roots to Confer Salt Tolerance. *Plant Physiol*, *184*(4), 2199-2215. https://doi.org/10.1104/pp.20.01054

Lauritano, C., Procaccini, G., & Ianora, A. (2012). Gene expression patterns and stress response in marine copepods. *Mar Environ Res*, *76*, 22-31. https://doi.org/10.1016/j.marenvres.2011.09.015

Lee, C. E., Charmantier, G., & Lorin-Nebel, C. (2022). Mechanisms of Na(+) uptake from freshwater habitats in animals. *Front Physiol*, *13*, 1006113. https://doi.org/10.3389/fphys.2022.1006113

Moya, A., Howes, E. L., Lacoue-Labarthe, T., Forêt, S., Hanna, B., Medina, M., Munday, P. L., Ong, J.-S., Teyssié, J.-L., Torda, G., Watson, S.-A., Miller, D. J., Bijma, J., & Gattuso, J.-P. (2016). Near-future pH conditions severely impact calcification, metabolism and the nervous system in the pteropod *Heliconoides inflatus*. *Global change biology*, *22*(12), 3888-3900. https://doi.org/https://doi.org/10.1111/gcb.13350

Parrish, K. K., & Wilson, D. F. (1978). Fecundity studies on *Acartia tonsa* (*Copepoda: Calanoida*) in standardized culture. *Marine Biology*, *46*(1), 65-81. https://doi.org/10.1007/bf00393822

Pizzagalli, M. D., Bensimon, A., & Superti-Furga, G. (2021). A guide to plasma membrane solute carrier proteins. *The FEBS Journal*, *288*(9), 2784-2835. https://doi.org/https://doi.org/10.1111/febs.15531

Podbielski, I., Hiebenthal, C., Hajati, M.-C., Bock, C., Bleich, M., & Melzner, F. (2022). Capacity for Cellular Osmoregulation Defines Critical Salinity of Marine Invertebrates at Low Salinity. *Frontiers in Marine Science*, *9*. https://doi.org/10.3389/fmars.2022.898364

Shayegan, M., Esmaeili Fereidouni, A., Agh, N., & Jani Khalili, K. (2016). Effects of salinity on egg and fecal pellet production, development and survival, adult sex ratio and total life span in the calanoid copepod, *Acartia tonsa*: a laboratory study. *Chinese Journal of Oceanology and Limnology*, *34*(4), 709-718. https://doi.org/10.1007/s00343-016-5030-4

Snyder, M. J. (2000). Cytochrome P450 enzymes in aquatic invertebrates: recent advances and future directions. *Aquat Toxicol*, *48*(4), 529-547. https://doi.org/10.1016/s0166-445x(00)00085-0

Sullivan, B. K., Costello, J. H., & van Keuren, D. (2007). Seasonality of the copepods *Acartia hudsonica* and *Acartia tonsa* in Narragansett Bay, RI, USA during a period of climate change. *Estuarine, Coastal and Shelf Science*, *73*(1-2), 259–267. https://doi.org/10.1016/j.ecss.2007.01.018

Tang, S., Wang, J., Yan, K., Yu, Z., & Li, J. (2025). Genome-Wide Identification of Solute Carrier Family 12 and Functional Characterization of Its Role in Saline-Alkaline Stress Acclimation in the Ridgetail White Shrimp *Exopalaemon carinicauda*. *Int J Mol Sci*, *26*(17). https://doi.org/10.3390/ijms26178339

Witalis, B., Iglikowska, A., Ronowicz, M., Weydmann-Zwolicka, A., & Kukliński, P. (2024). The zooplankton community of Baltic Sea ports: diversity and seasonal dynamics. *Oceanologia*. https://doi.org/10.1016/j.oceano.2024.02.001

Yang, Z., Zhou, J., Zhu, L., Chen, A., & Cheng, Y. (2022). Label-free quantification proteomics analysis reveals acute hyper-osmotic responsive proteins in the gills of Chinese mitten crab (*Eriocheir sinensis*). *Comp Biochem Physiol Part D Genomics Proteomics*, *43*, 101009. https://doi.org/10.1016/j.cbd.2022.101009
